# IFNAR1⁺ Neutrophils Orchestrate Chronic Inflammatory Damage Through Mitochondrial Remodeling

**DOI:** 10.1101/2025.09.22.677957

**Authors:** Cecilia Pessoa Rodrigues, Raquel R. Calçada, Mafalda Arnaud, Lukas Kaltenbach, Mika Manser, Anja Scheidegger, Leon Sidney Strauss, Alina Gavrilov, Sharang Kulkarni, Ece Yildiz, Daniela Liberati, Mairene Coto-Llerena, Tamara Hoening, Markus Martin Kramberg, Julian Behr, Anna Mechling, Marvin Hering, Kara G. Lassen, Oliver Soehnlein, Emma Doran, Daniel Regan-Komito

**Author notes:** Senior author (Emma Doran, Daniel Regan-Komito).

## Abstract

Neutrophils are abundant innate effector cells that drive mucosal inflammation, yet the mechanisms by which they contribute to chronic inflammatory diseases across distinct tissues remain incompletely understood. Here, by reanalyzing single-cell RNA-seq datasets from patients with inflammatory bowel disease (IBD) and chronic obstructive pulmonary disease (COPD), we identify a shared neutrophil activation program enriched for type I interferon (IFN) signaling, nuclear factor-κB (NF-κB) and AP-1 transcriptional regulators, and effector pathways including NETosis, degranulation, and leukocyte trafficking. To interrogate these signatures, we established a CRISPR-compatible neutrophil differentiation platform from adult CD34⁺ progenitors, which yielded cells closely resembling primary neutrophils at transcriptomic, proteomic, and functional levels. A targeted CRISPR-Cas9 screen revealed a central role for the mitochondrial iron transporter mitoferrin-1 (SLC25A37) in coordinating neutrophil oxidative phosphorylation, NET formation, and type I IFN production downstream of TLR9. Mechanistically, we show that NET-derived citrullinated histones activate an autocrine IFNα–IFNAR1 loop, amplifying neutrophil inflammatory functions without impairing phagocytosis. Disruption of this loop, through IFNAR1 depletion or blockade, dampened neutrophil-driven tissue damage in human intestinal and alveolar organoid co-cultures as well as in murine models of colitis and cigarette smoke–induced lung inflammation. These findings uncover a conserved IFN-driven metabolic circuit in neutrophils that underpins pathology across chronic mucosal diseases and identify IFNAR1 as a therapeutic node to selectively disarm neutrophil-mediated tissue injury.

## Introduction

Neutrophils, the most abundant white blood cell in circulation, are essential for the innate immune defence due to their rapid response to infections and tissue damage. Neutrophils neutralize pathogens using several key mechanisms, including phagocytosis, degranulation, the release of neutrophil extracellular traps (NETosis), and the production of reactive oxygen species (ROS)^1,2^. These functions, under normal circumstances, are self-limiting^3^, crucial for host defense and maintaining homeostasis by initiating the reparative process in mucosal environments like the gastrointestinal and respiratory tract^4^.

While neutrophils play a vital role in clearing pathogens, their excessive accumulation and sustained activation during chronic inflammation can compromise epithelial barrier integrity, amplify inflammatory signaling, and drive persistent immune activation^5–9^. In chronic inflammation, neutrophils are continuously recruited and exacerbate tissue damage by releasing mediators such as serine proteases, cytokines—which activate other immune cells—and ROS, thereby sustaining the inflammatory milieu^10,11^. In addition, increased neutrophil extracellular traps (NETs) correlates with active diseases in chronic mucosal inflammatory diseases^8,12^.

This maladaptive response contributes significantly to the chronic tissue injury observed in chronic mucosal inflammation and diseases, including, inflammatory bowel disease (IBD), encompassing Crohn’s disease (CD) and ulcerative colitis (UC)^12,13,14^. A similar pathogenic paradigm is observed in chronic inflammatory airway diseases such as chronic obstructive pulmonary disease (COPD), where neutrophilic inflammation is a defining feature. In COPD, neutrophil infiltration promotes airway remodeling, mucus hypersecretion, and progressive alveolar destruction (emphysema)^15^. These outcomes underscore the central role of neutrophil-driven inflammation in shaping both disease severity and progression^16,17,9^. Therefore, understanding the mechanisms that control neutrophil activation, migration, and effector functions is critical for the development of successful strategies to reduce their harmful effects while preserving their essential immune functions.

Using human IBD and COPD diseases data, we identified the crucial role of mitoferrin-1 (*SLC25A37*) in mediating neutrophil activation. We found that the mitochondrial iron uptake functions as a rheostat that promotes increased mitochondrial mass activity due to high fusion and mitochondrial oxygen consumption. We observed that mitochondrial dynamics are different for extra and intracellular insults in which mitochondrial fusion and oxygen consumption are required for the type I IFN production by neutrophils. IFN-α then engages with IFNAR1 on neutrophils, reinforcing a positive feedforward loop that sustains the inflammatory response.

Using the IBD and COPD mouse models and 3D-organoid cocultures we report that anti-IFNAR1 treatment inhibited neutrophil inflammation and tissue damage. These findings can pave the way for innovative approaches aiming at modulating neutrophil activity in mucosal immunology.

## Results

### Infiltrating neutrophils share signatures in COPD and IBD

To identify shared neutrophil pathway modulations in IBD gut tissue and COPD lungs, we curated and analyzed 15 publicly available scRNA-seq datasets (7 IBD and 8 COPD) from patients and healthy controls. After selecting CD45+ (*PTPRC* expressing cells) leukocytes, we used the CellTypist tool to standardize neutrophil annotations across datasets (Fig. 1a). Pseudo-bulk analysis comparing patient and healthy samples revealed both disease-specific and shared neutrophil upregulated genes (Fig. 1b, Extended Data Table 1).

**Figure 1.**
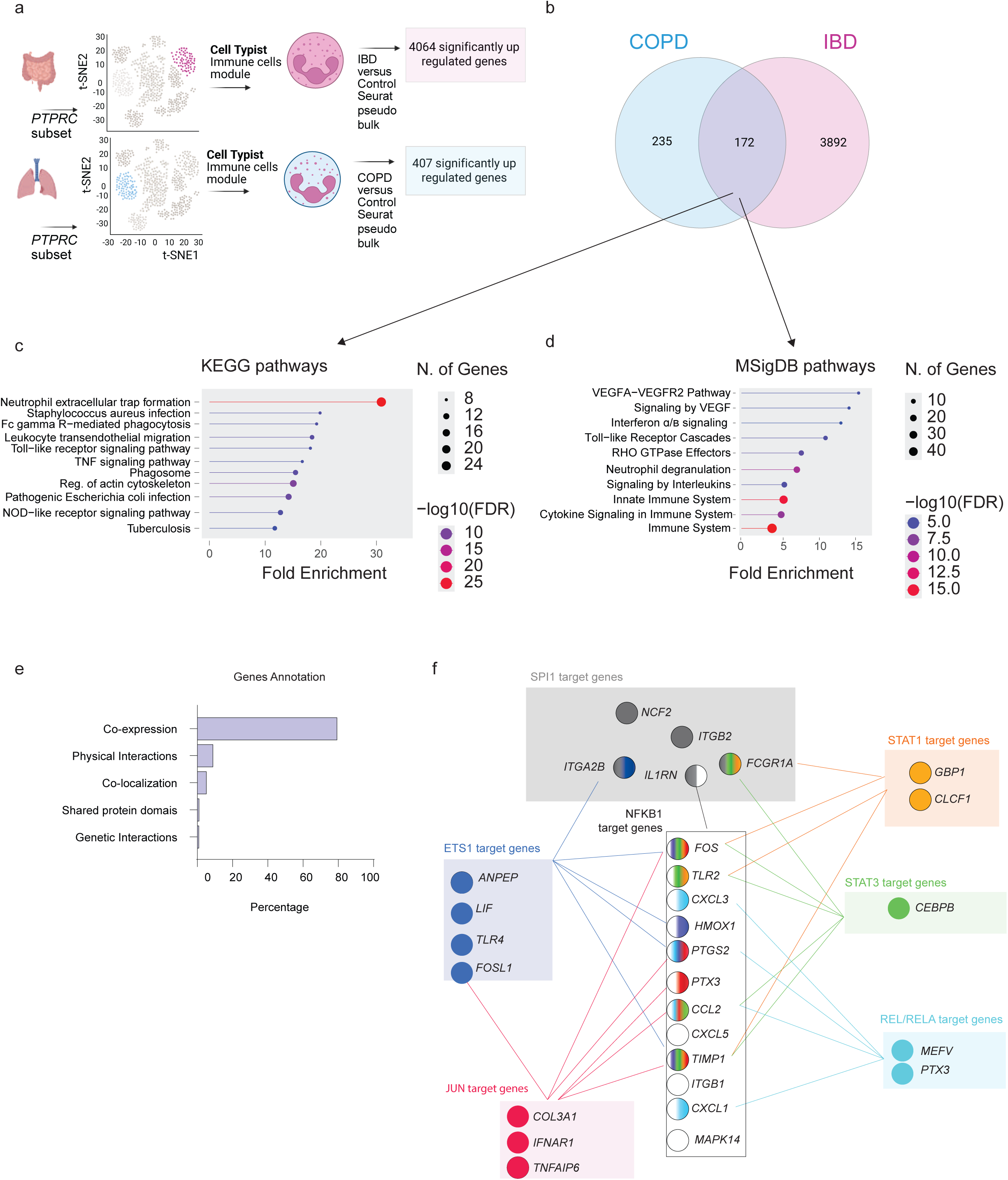
Characterization of upregulated genes and pathways found in diseased infiltrating neutrophils. a) Graphical scheme depicting the scRNAseq strategy to harmonize the neutrophil infiltration profile in IBD and COPD. b) Venn diagram showing the disease-specific and common up-regulated genes. c) KEGG pathways and d) MSidDB hallmarks associated with the shared upregulated genes in COPD/IBD infiltrating neutrophils. e) Description of the expression pattern from the shared upregulated genes. The software “geneMania” was used for the classification. f) Transcription factor network analysis (TF-TRED) showing the active circuits based on the shared upregulated genes in COPD/IBD infiltrating neutrophils. The TF pathways are depicted by colors and the genes belonging to the TF network are labeled with the colors. Genes belonging to more than one TF are connected with a line and the respective TF color.

Shared upregulated genes highlighted key neutrophil functions such as NET formation, leukocyte migration, degranulation, and interleukin signaling^18^ (Fig. 1c-d). Notably, we observed strong enrichment of type I interferon (IFN) pathways (Extended Data Fig. 1a-e). These have been previously implicated in Dextran sulfate sodium (DSS)-colitis and IBD patient samples and linked to genetic variants like the IFNAR1-associated SNP rs2284553^19,20^. In COPD, IFN signaling is similarly relevant, with elevated *ISG15* levels during exacerbations^18^.

Shared genes exhibited strong co-expression (Fig. 1e), and transcription factor analysis revealed activation of inflammatory regulators including *RELA, NFKB1, REL, JUN, SPI1, STAT1, and ETS1* (Fig. 1f). These results point to a common neutrophil-driven inflammatory program across IBD and COPD, particularly involving type I IFN signaling.

### CD34-Adult derived neutrophils displayed similar phenotype as fresh isolated neutrophils

To investigate the function of shared upregulated genes in IBD and COPD neutrophils, we developed a CRISPR-compatible in vitro neutrophil differentiation model from human CD34+ hematopoietic stem and progenitor cells (HSPCs) sourced from adult (AD) or cord blood (CB) (Fig. 2a). The three-stage protocol involved: (1) expansion of the progenitor cells using the OCI-AML22 cytokine cocktail^21^, (2) myeloblast lineage specification, and (3) neutrophil differentiation using a modified cytokine mix. Notably, rosiglitazone, a PPARɣ agonist, was added to enhance differentiation.

**Figure 2.**
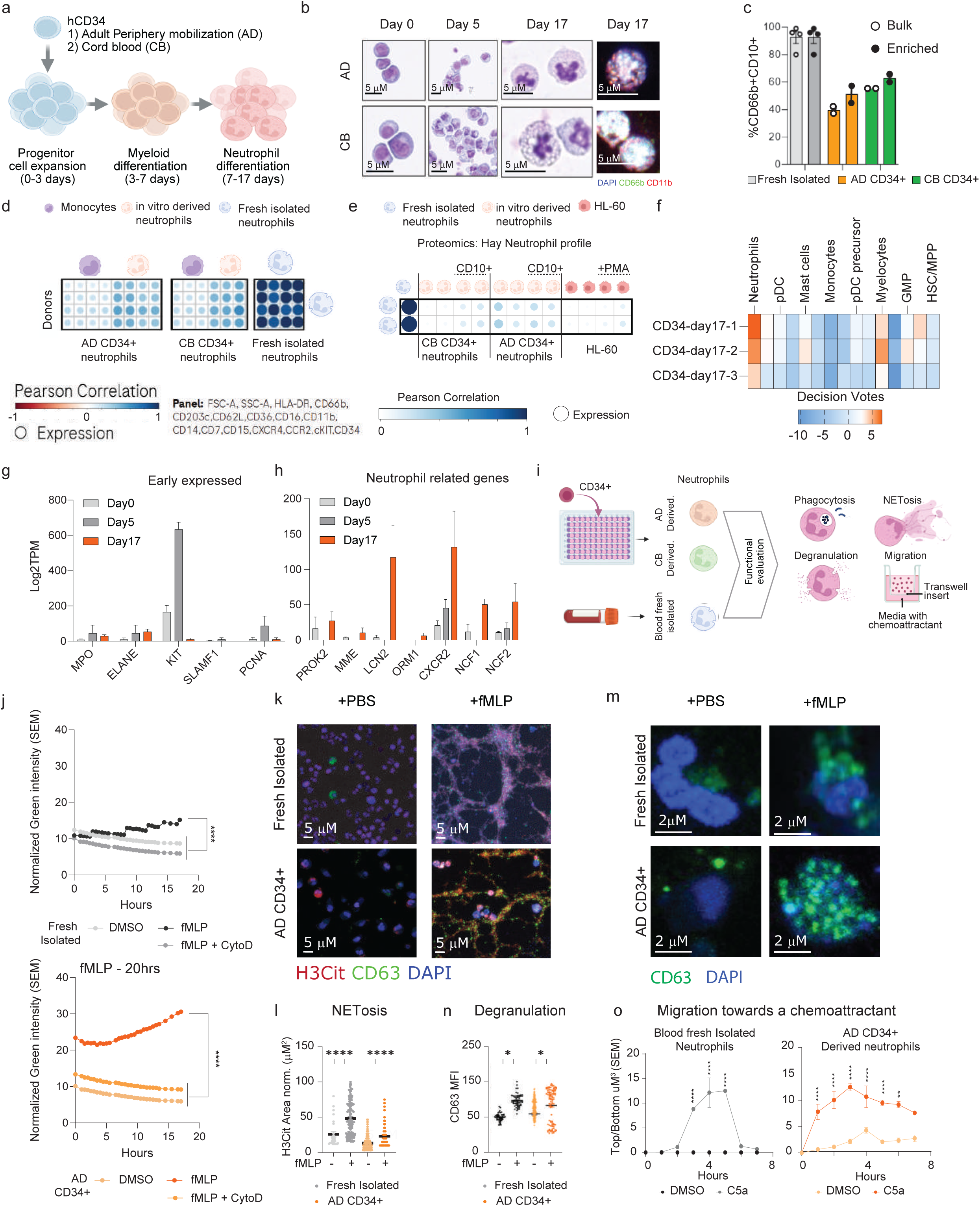
Phenotypical and functional characterization of the in vitro derived neutrophils. a) Schematic showing the in vitro strategy to derive neutrophils from CD34+ cells. b) Representative hematoxylin and eosin staining images (n=2) across the neutrophil differentiation process. c) Quantification of CD11b+CD66b+ cells before (bulk) or after CD10+ enrichment (enriched). d) High dimensionality flow cytometry analysis comparing fresh isolated neutrophils (blue cells) with CD34-derived neutrophils (orange) or monocytes (purple) (n=4). Pearson correlation is depicted by the heatmap key and the circle size represent the MFI levels for HLA-DR, CD66b, CD203e, CD62L, CD36, CD16, CD11b, CD14, CD7, CD15, CXCR4, CCR2, cKit and CD34. e) Correlation plot showing the expression and Pearson correlation of fresh isolated neutrophils, in vitro derived neutrophils plus or minus CD10+ further enrichment. We used the Hay Neutrophil profile as a reference to subset the relevant proteins from the proteomics from the MSigDB. f) Celltypist unbias voting for 3 independent in vitro differentiation cultures showing high decision votes to the neutrophil profile. g) Progenitor cell gene or h) mature neutrophil gene expression across the in vitro derived neutrophil differentiation culture. i) Schematic showing the functions tested in AD-CD34+ derived neutrophils and fresh isolated neutrophils. j) E-Coli ph-Rhodo incorporation across 18 hours live imagining. Neutrophils were activated with fMLP and cytochalasin D was used as a negative control. The phagocytosis capacity was measured by the green channel intensity. Data was normalized by the total number of cells present in the field of view (n=3 independent experiments; 3 replicates per assay). Statistical analysis was performed by two-way ANOVA followed by Sidak’s multiple comparison. p***<0.001. k) Representative images showing NETs, TRAP formation, upon fMLP treatment. NETosis is depicted by histone 3 citrullination (H3Cit, red), nuclei (DAPI, blue) and degranulation is depicted by CD63 (green). l) Quantification of the TRAP area. Statistical analysis was performed by one-way ANOVA followed by Tukey’s multiple comparison. p****<0.0001. m) Representative images showing neutrophil degranulation depicted by CD63 (Green) upon fMLP treatment. Nuclei is depicted by DAPI (blue). n) Quantification of the CD63 MFI. Statistical analysis was performed by one-way ANOVA followed by Dunn’s multiple comparison. p*<0.05. o) Line plots showing the migration capacity towards C5 (chemoattractant) across 8 hours live imaging. (n=3 independent experiments; 3 replicates per assay), Statistical analysis was performed by two-way ANOVA followed by Sidak’s multiple comparison. p****<0.0001. Data variation is presented as ±SEM.

After 17 days, ∼60-80% of cells showed neutrophil-like morphology, with increased CD66b+ and decreased CD34+ populations (Fig. 2b and Extended Data Fig. 2a), indicating granulocyte lineage commitment. Neutrophils were isolated via negative selection or by negative selection followed by positive selection of CD10+ cells, which did not trigger activation (Extended Data Fig. 2b). Up to 60% mature neutrophils (CD66b+CD10+) were confirmed by cytospin and flow cytometry (Fig. 2b-c), with no contamination from other mature immune cells.

High-dimensional flow cytometry revealed ∼70% similarity between AD and CB in vitro derived and fresh neutrophils, versus ∼10% similarity with monocytes (Fig. 2d). Proteomic profiling showed AD-derived neutrophils more closely resembled fresh cells, while CB-derived and HL-60 cells clustered apart (Fig. 2e, Extended Data Fig. 2c). Pathway analysis highlighted stronger immune responses and increased protein translation in AD-derived neutrophils (Extended Data Fig. 2d-e). Collectively, suggesting that at the protein level the AD-derived neutrophils retain fresh isolated neutrophils similarities, while HL-60 and CB-derived showed lower similarity.

Bulk RNA-seq across the differentiation timeline confirmed a transition from progenitor to mature neutrophil signatures by day 17 (Fig. 2f-2h, Extended Data Fig. 2f). Overall, along with the proteomics, AD-derived neutrophils best mimic fresh neutrophils at transcriptomic levels.

To evaluate the functionality of in vitro-derived neutrophils, we tested phagocytosis, NETosis, degranulation, and migration (Fig. 2i, Extended Data Fig. 2g). The in vitro derived neutrophils displayed enhanced phagocytic capacity (Fig. 2j, Extended Data Fig. 2h), consistent with reports of higher phagocytic activity in cord blood monocytes^22^. NETosis analysis revealed that CB-derived neutrophils released histones but failed to form extracellular traps, whereas AD-derived cells formed structurally normal, though smaller, traps a phenotype similar to fresh neutrophils (Fig. 2k-l, Extended Data Fig. 2i). In degranulation assays, in vitro-derived neutrophils responded comparably to fresh cells upon fMLP stimulation (Fig. 2m-n, Extended Data Fig. 2j). For migration, in vitro-derived neutrophils responded to C5a more rapidly, initiating movement within 1 hour versus 3 hours for fresh cells (Fig. 2o, Extended Data Fig. 2k). Collectively, confirming that AD-derived neutrophils resemble fresh isolated neutrophils.

Further testing confirmed that AD-derived neutrophils responded to LPS and PMA. Upon LPS exposure, they increased IL-1β and IFN-γ, decreased IL-10 and CD62L, and upregulated CXCR2 (Extended Data Fig. 2l-p), indicating an activated phenotype. Overall, AD-derived neutrophils closely mimic the functional behavior of fresh neutrophils, supporting their use in downstream perturbation studies.

### Mitoferrin-1 regulates neutrophil mitochondrial dynamics and metabolic responses to TLR agonists

To investigate how genes upregulated in infiltrating neutrophils from IBD and COPD patients (Fig. 1b) contributed to neutrophil dysfunction, we performed an array CRISPR-Cas9 screen. We prioritized candidate genes whose proteins were significantly enriched in neutrophils based on the human protein atlas (Fig. 3a and Extended Data Table 2). We then employed a multiplexed readout to assess their contribution to key neutrophil functions (Fig. 3b).

**Figure 3.**
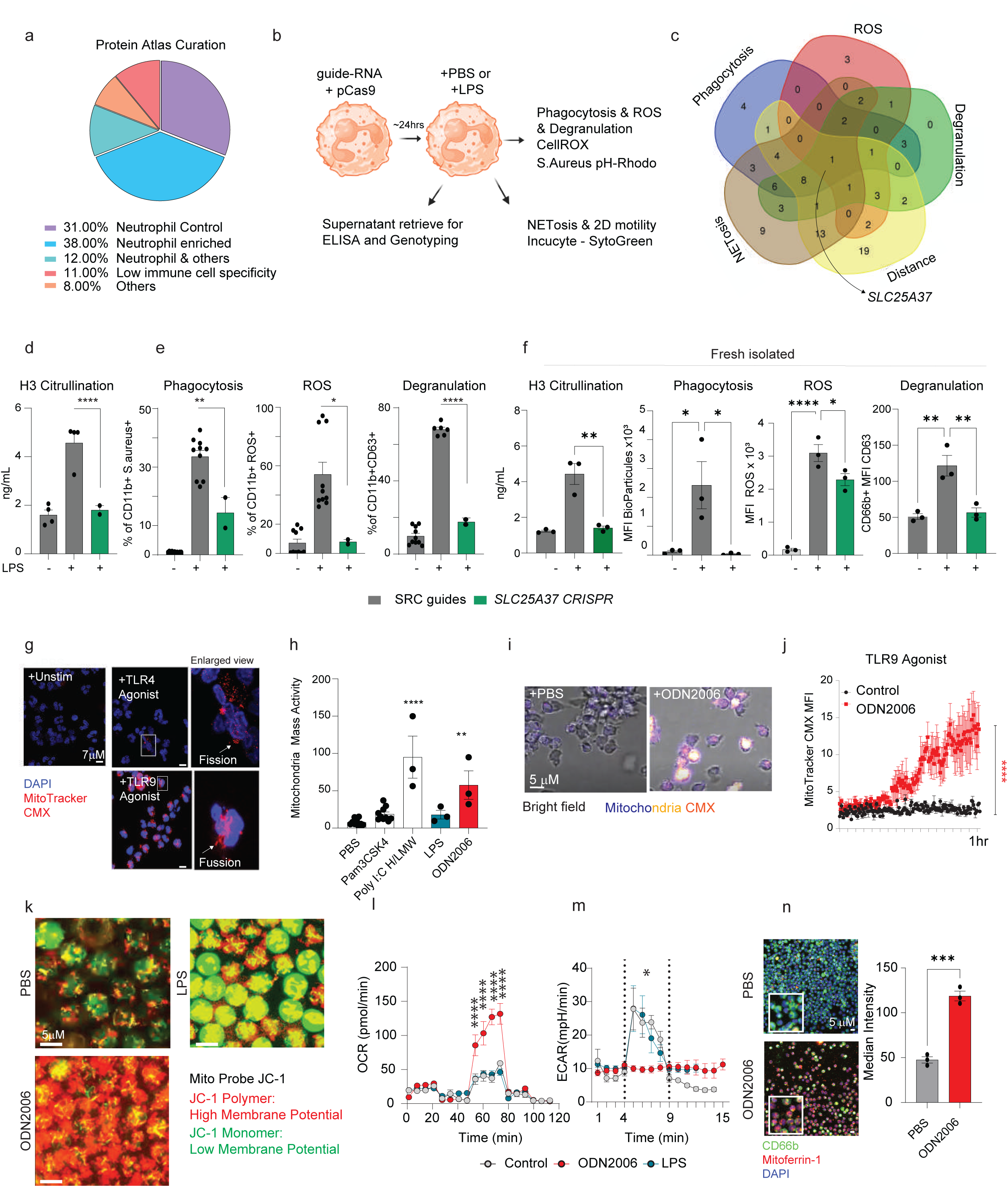
Mitochondrial dynamics are required for neutrophil activation. a) Annotation of the 95 targets identified in the scRNAseq analysis. b) Graphic scheme showing the multifunctional readouts employed for the CRISPR-cas9 screen. c) Venn diagram depicting the 31 functional models and the number of targets affecting the functions. Only significant (P<0.05) targets are shown in the diagram. d) Bar plots showing the histone 3 citrullination present in the supernatant. LPS was used as positive control and cl-Amidine, a PAD4 inhibitor, was used as negative control. The complete number of targets significantly affecting NETosis is shown in Extended Data Fig. 4. e) Barplots showing the phagocytosis, ROS and degranulation capacity of *SLC35A37* depleted in vitro derived neutrophils (from the CRISPR-Cas9 screen; n=3) and f) as e) but from fresh isolated neutrophils. Statistical analysis was performed by Kruskall Wallis test followed by Dunnett’s multiple comparison. p*<0.05, p**<0.01,p***<0.001 and p****<0.0001. The CRISPR-Cas9 screen was performed twice in independent differentiation experiments. g) Representative immunofluorescence showing the mitochondrial mass activity, red-CMX mitochondrial staining, in steady state (PBS) and stimulated with an extracellular insult, TLR4 agonist (LPS) or an intracellular insult, TLR9 agonist (ODN2006). The nucleus was stained using DAPI. h) Quantification of the mitochondrial mass activity under steady state (n=10) or upon extracellular insults TLR2, Pam3CSK4 (n=10), TLR4, LPS (n=3) or intracellular insults TLR3, high molecular weight and low molecular Poly I:C (n=3) and TLR9, ODN2006 (n=3). The dots in the plot represent a donor. Statistical analysis was performed by one-way ANOVA followed by Dunnett’s multiple comparison. p**<0.01 and p****<0.0001. Samples were acquired 4 hours after TLR agonist and Golgi-block stimulation. i) Representative image from a time lapse live imaging assay showing the mitochondrial mass activity in steady state (PBS) and TLR9 treated cells. j) Quantification of i) over 1 hour imaging (n=3). Statistical analysis was performed by two-way ANOVA. p****<0.0001. k) Representative image showing the JC-1 mito probe staining. Green color represents depolarized mitochondria and red represents polarized mitochondria. The quantification (n=3) is shown in Extended Data Fig. 4. l) Real-time cell metabolic analysis (Seahorse) showing the oxygen consumption capacity (OCR) and m) extracellular acidification rate (ECAR) in steady state or treated with TLR9 agonist (ODN2006) or TLR4 agonist (LPS) fresh isolated neutrophils (n=5).Statistical analysis was performed by two-way ANOVA. p****<0.0001. n) Left panel: Representative image from 3 donors showing the levels of *SLC25A37* (mitoferrin-1) under steady state or after ODN2006 treatment. Right panel: Cellular median intensity quantification of SLC25A37 (mitoferrin-1) levels. Normal distribution was evaluated by the Shapiro Wilk test followed by the Student-t test. ***p=0.0004. Data variation is presented as ±SEM.

Initially, we optimized the transfection efficiency of guide RNAs derived from the GeCKO library^23^ (Extended Data Fig. 3a-d). We then conducted a targeted screen of the 95 genes, evaluating neutrophil activity through multiple functional assays, namely phagocytosis, ROS production, degranulation, two-dimensional (2D) motility (“distance”), and NET formation (NETosis) (Extended Data Fig. 3e-g). Notably, only viable cells were considered for the phagocytosis, ROS and degranulation readouts.

To visually convey the intricate relationships among the five assessed functional parameters, we employed a Venn diagram that highlights their overlapping nature. Focusing our analysis on statistically significant alterations relative to LPS-pertubation (Fig. 3c and Extended Data Fig. 3g; Extended Data Table 2). This strategic approach enabled the delineation of discrete functional modalities across the explored readouts.

For instance, our assay uncovered six candidate genes - one of which is *ISG15*, whose perturbation appeared to compromise degranulation, NETosis and phagocytosis (Extended Data Table 2). Unveiling a compelling link between type I interferon signaling and neutrophil functional capacity. We then focused on the candidate gene that impaired all the five explored functions, *SLC25A37* (mitoferrin-1). In our screen, we identified that *SLC25A37* plays a central role in regulating all the neutrophil functions assessed (Fig. 3d-e), with similar findings observed in viable freshly isolated neutrophils (Fig. 3f, Extended Data Fig. 3h-k).

Mitoferrin-1 is a mitochondrial protein controlling the balance of iron in the cytoplasm and mitochondria^24^, which is required for proper mitochondria oxidative phosphorylation and dynamics^25^. Mitochondrial dynamics, either fission or fusion, has a crucial role in mediating innate immune responses^26^. Fission facilitates mitophagy and redistribution, while fusion maintains integrity and optimizes energy production. Disrupting this balance impacts immune activation, metabolism, and inflammation, linking mitochondrial dysfunction to autoimmune and chronic inflammatory diseases^27,28,29,30,31^. Then, given the importance of toll-like receptors (TLR) in mediating neutrophilic chronic mucosal inflammation^32^ and the role of mitoferin-1 in maintaining mitochondria homeostasis^24^, we sought to investigate the mitochondrial dynamics, fusion and fission, in fresh isolated neutrophils in response to different TLR-agonists.

By mitochondrial imaging we found that while TLR2 (Pam3CSK4) and TLR4 (LPS) promoted a mitochondrial fission like behavior, TLR3 (Poly I:C, high plus low) and TLR9 (ODN2006) promoted fusion (Fig. 3g-h and Extended Data Fig. 3l). Suggesting that extracellular or intracellular pathogen associated patterns may elicit a distinct mitochondrial program.

Among the explored TLRs, TLR9 ligation produced the fastest response time of mitochondrial mass activity (Fig. 3j, Extended Data Fig. 4a-b, Extended Data Video1 and 2) and neutrophil activation (Extended Data Fig. 4c-e), with hyperpolarization of the mitochondria, unlike LPS (Fig. 3k). To understand if these observations occur due to mitochondrial collapse or the iron uptake mediated by mitoferrin-1, we treated the cells with the iron chelator, deferasirox (DFX)^33^. DFX treatment blocked the mitochondrial polarization upon TLR9 agonist challenge, suggesting that iron uptake is crucial for mitochondrial polarization in neutrophils (Extended Data Fig. 4f). This indicates that the mitochondrial iron uptake upon TLR9 stimulation may act as a driver for mitochondrial polarization and fusion.

Next, we performed real-life metabolic rate analysis by Seahorse extracellular flux analysis, and found that the oxygen consumption rates (OCR) were significantly higher in TLR9-stimulated neutrophils (Fig. 3l), whereas LPS led to increased glycolysis (Fig. 3m). Mitotracker-Green staining confirmed mitochondrial morphology changes upon TLR4 or TLR9 stimulation in fresh neutrophils (Extended Data Fig. 4g). Moreover, we observed that TLR9 agonism promotes the expression of mitoferrin-1 (Fig. 3n) and increased mitochondrial area, an effect absent in mitoferrin-1 depleted-fresh isolated neutrophils (Extended Data Fig. 4h). This suggests that the observed changes in response to TLR9 ligand (ODN2006) are driven by mitochondrial dynamics, which promote oxidative phosphorylation upon intracellular insults, such as a virus infection. However, infection from extracellular bacteria, modelled here by TLR4 ligand (LPS) and TLR2 ligand (Pam3CSK4) challenge, induce glycolysis.

In the context of mucosal inflammation, it is notable that we observed enrichment of type I IFN and virus-like response in the infiltrating neutrophils (Fig. 1, Extended Data Fig. 1c-d), suggesting that, in a disease context, neutrophils may display increased OxPhos and mitochondrial activity. Altogether, our data indicate that different insults could elicit a distinct mitochondrial morphology and membrane potential in neutrophils.

### Mitochondrial mass activity regulates type I Interferon production in neutrophils

To further understand the role of *SLC25A37* in regulating the type I IFN-driven neutrophil mitochondrial behaviour we transfected fresh isolated neutrophils with CRISPR-Cas9 guides targeting *SLC25A37* (mitoferrin-1) (Extended Data Fig. 3h-i). We found that *SLC25A37* depleted-neutrophils displayed lower OCR (Fig. 4a) and did not respond to TLR9 ligand (Fig. 4b). These results support the decreased NETosis capacity observed in the *SLC25A37* and *OPA1* depleted AD-neutrophils (Fig. 3c, Extended Data Table 2). Collectively, the data suggests that mitoferrin-1 plays a crucial role in mediating neutrophil metabolic responses upon TLR9 stimulation.

**Figure 4.**
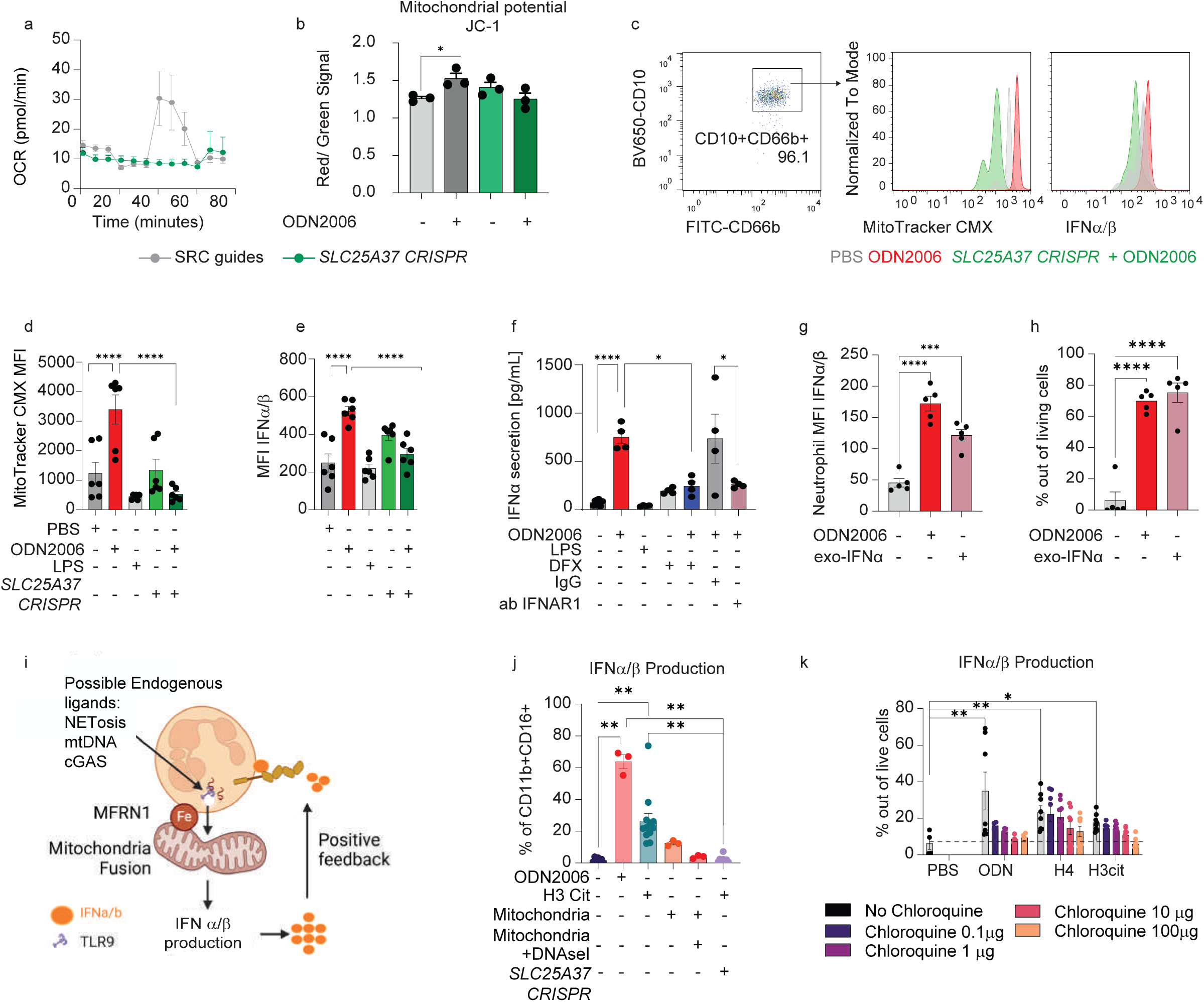
Mitochondria activation is required for IFN-ɑ production. a) Real-time cell metabolic analysis (Seahorse) showing the oxygen consumption rate (OCR) of control neutrophils (transfected with a scramble gRNA, grey, n=3) and *SLC35A37/*mitoferrin-1 depleted neutrophils (green, n=3). b) Quantification of the mitochondria polarization based on JC-1 mito probe red versus green ratio. Green color represents depolarized mitochondria and red represents polarized mitochondria (n=3). Control neutrophils (transfected with a scramble gRNA, black) and *SLC35A37/*mitoferrin-1 depleted neutrophils (green). c) Representative flow cytometry histogram depicting the mitochondrial mass activity (left panel) and type I IFN levels. ODN2006, the TLR9 agonist was used as positive control (red line). d) Quantification of the mitochondrial mass activity and e) type I IFN levels, PBS (n=6), ODN2006 (n=6), LPS (n=5), *SLC35A37*/mitoferrin-1 depleted (n=5, n=6). The “+” under the bar represents the addition of the compound. f) ELISA quantification showing the amount of IFN-ɑ present in the supernatant of stimulated neutrophils. The “+” under the bar represents the addition of the compound. DFX: Deferasirox. IgG: Isotype IgG control. Ab: Antibody. g) Quantification of IFN-ɑ levels and h) percentage of IFN-ɑ positive neutrophils upon PBS, ODN2006 or exogenous IFN-ɑ challenged (n=5). MFI: Mean Fluorescent Intensity. Statistical analysis was performed by one-way ANOVA followed by Tukey’s multiple comparison. Data variation is presented as ±SEM. i) Schematic representation of the proposed positive feedback loop exerted by IFN-ɑ and its receptor IFNAR1. j) Bar plot showing the percentage of IFN-ɑ producing neutrophils upon PBS (steady state control, n=7), ODN2006 (n=3), Histone 3 citrullination (H3) (n=10), autologous Mitochondria (n=3), autologous Mitochondria plus DNAseI (n=3), and *SLC35A27*/mitoferrin-1 depleted neutrophils (n=7). Besides the *SLC35A27*/mitoferrin-1 depleted group the other groups were transfected with the scramble gRNA control. Statistical analysis was performed by Kruskal-Wallist test followed by Dunn’s multiple comparison. **p<0.01. k) Quantification of type I IFN production upon ODN2006 and NETosis protein (histone 4, H4, and Histone 3 citrullination, H3Cit). The cells were co-treated with chloroquine in order to test the endosomal formation dependence of the response.

To investigate the role of mitochondrial mass activity in neutrophil-mediated immune responses, we stimulated cells with the TLR9 agonist ODN2006, which significantly increased mitochondrial activity and type I IFN production (Fig. 4c-e). This effect was absent in *SLC25A37* CRISPR neutrophils (Fig. 4c-e), suggesting that mitoferrin-1 is required for this response. A positive correlation between mitochondrial mass activity and type I IFN production further supports this relationship (Extended Data Fig. 4i). Additionally, treatment with DFX blocked type I IFN production (Extended Data Fig. 4j). To further validate these findings, we measured IFN-α secretion in neutrophil supernatants following TLR9 stimulation. Consistent with our previous results, DFX treatment significantly impaired IFN-α secretion, reinforcing the link between mitochondrial metabolism and interferon signaling (Fig. 4f).

Because we have identified type I IFN signalling as a shared upregulated circuit in IBD and COPD infiltrating neutrophils (Fig. 1d-f; Extended Data Fig. 1a-e), we hypothesized that secreted IFN-α could contribute to a positive feedback loop, amplifying neutrophil activation. To test this, we stimulated the neutrophils with exogenous IFN-α in the presence or absence of an IFNAR1-neutralizing antibody. We also added a protein transporter inhibitor containing monensin to inhibit Golgi function to ensure that the paracrine/autocrine IFN-α is not involved (Fig. 4f-h). It is important to note that other inflammatory cytokines like IL-8 remained unchanged upon IFNAR1 antibody treatment (Extended Data Fig. 4k), while it decreased their mitochondrial mass activity upon TLR9 stimulation (Extended Data Fig. 4l). These experiments collectively suggest that IFN-α signaling may sustain an autocrine loop in neutrophils in response to TLR-9 stimulation.

Next, we sought to identify the endogenous ligand that could trigger the IFN-α/IFNAR1 axis in IBD and COPD to initiate the positive feedback mechanism in situ (Fig. 4i). Considering the DNA nature of ODN2006, we hypothesized that the DNA present in situ could act as the endogenous trigger, as shown previously in COPD^34^. We explored if autologous mitochondrial DNA or NETosis products (H3Cit) could promote IFN-α production. H3Cit, but not mitochondrial DNA, facilitated IFN-α production (Fig. 4j and Extended Data Fig. 4m-n). Corroborating with the role of mitoferrin-1 in regulating the IFN-α production, *SLC25A37* depletion abrogated the H3Cit response (Fig. 4j). Next, we sought to investigate the molecular mechanisms that may be responsible for the IFN-α production. As expected, when neutrophils were treated with ODN2006 and a pan-TLR inhibitor (MyD88i) there was a significant decrease in IFN-α production (Extended Data Fig. 4o). Contrasting to this, when we treated the cells with a cGAS-STING inhibitor we did not observe a decrease in IFN-α levels (Extended Data Fig. 4i), suggesting that TLR9 was the main sensor facilitating IFN-α production.

Because the MyD88i impaired the IFN-ɑ production, we hypothesized that neutrophils sensed DNA through TLR9. We therefore examined whether neutrophil-TLR9 response is involved in the recognition of NETosis components (H4 and H3Cit). IFN-ɑ induced by H4 and H3Cit was potently inhibited by chloroquine in a dose dependent manner (Fig. 4k), which blocks endosomal TLR signalling^35^. This suggests that NETosis may trigger TLR9 response to promote IFN-α production that can be further amplified by the positive feedback loop between IFN-α and IFNAR1. Indeed, a subset of neutrophils have been shown to express a set of interferon stimulated genes (ISGs) including *Ifit3* and *Isg15*^36,37^. This subset of neutrophils significantly increased in *E.Coli*-challenged hosts, suggesting an innate memory phenotype of these cells^36,37^. Here, we have added another layer to this population by highlighting a potential link between IFNAR1^+^ infiltrating neutrophils and inflammatory disease pathogenesis.

### Therapeutic IFNAR1 blockage ameliorates neutrophil-driven inflammation in models of IBD and COPD

To further explore the implications of IFNAR1 and the positive feedback loop of type I IFN and its receptor on IBD and COPD inflammation, we investigated whether IFNAR1 treatment could ameliorate the inflammatory milieu and disease activity (Fig. 5 and Extended Data Fig. 5).

**Figure 5.**
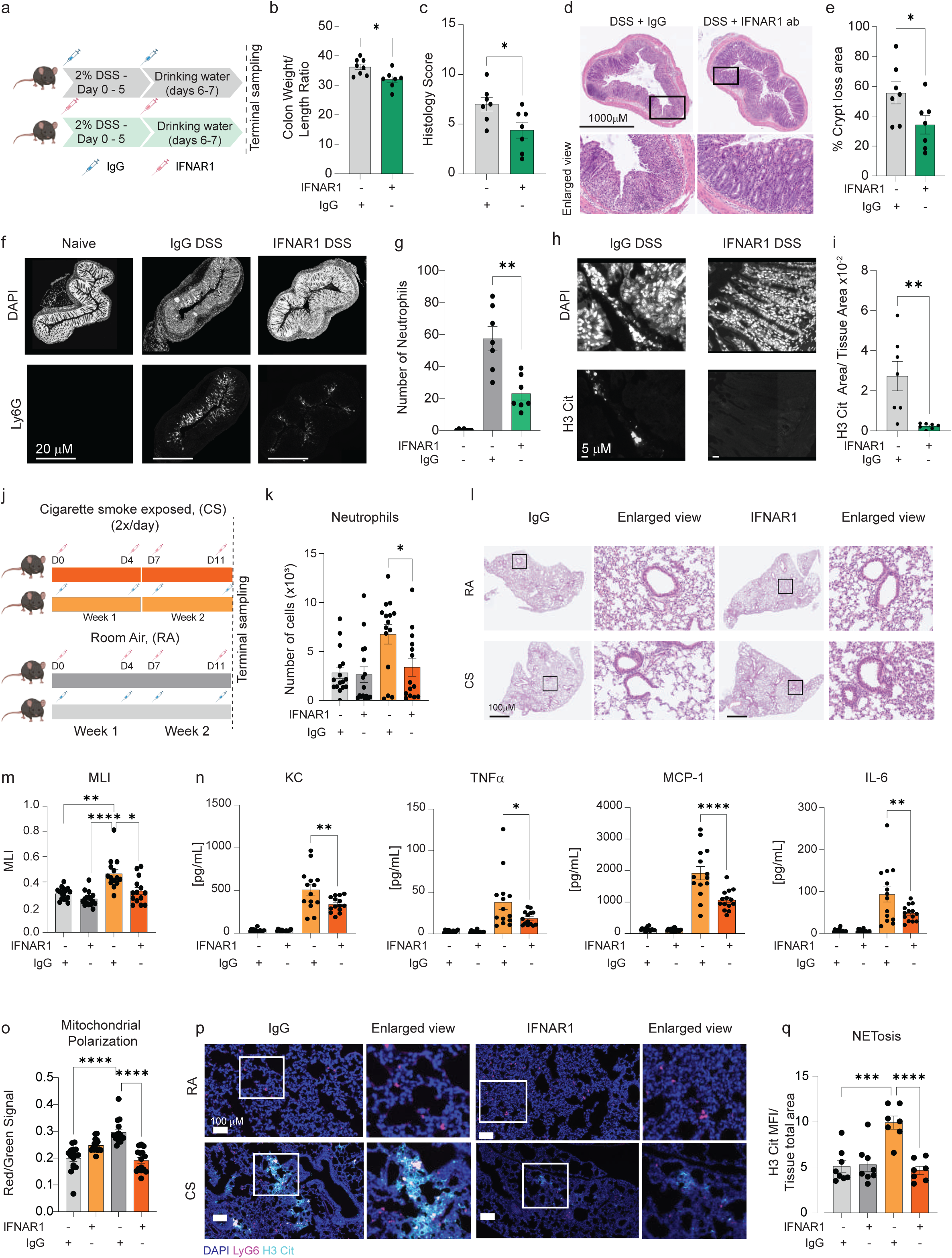
Pharmacological inhibition of the IFNAR1 signalling affected neutrophil-pathogenic functions in vivo. a) Graphic scheme outlining the DSS-Colitis model with the anti-IFNAR1 or IgG treatment. b) Colon weight (g) over length (cm) ratio. Statistical analysis was performed by Student t-test after passing data normality. c) Total histology score based on leukocyte infiltration, tissue damage and ulceration. Statistical analysis was performed by Student t-test after passing data normality. d) Hematoxylin and eosin representative staining of the colon. The enlarged view shows the crypt status. e) Quantification of the crypt integrity. f) Representative imaging showing the neutrophils infiltration depicted by Ly6G staining in the colon. The nucleus was stained with DAPI. g) Quantification of the number of Neutrophils observed in the colon. Data passed the Shapiro-Wilk test normality and statistical analysis was performed by One-way ANOVA followed by Tukey’s multiple comparison. h) Representative imaging showing H3 citrullination (H3 Cit) in situ. i) Quantification of the number of H3 Cit. Data passed the Shapiro-Wilk test normality and statistical analysis was performed by One-way ANOVA followed by Tukey’s multiple comparison. j) Graphic scheme showing the cigarette smoked COPD model with the anti-IFNAR1 or IgG treatment. k) Quantification of the infiltrating neutrophils in RA and CS animals treated or not with IFNAR1 antibody (500 ug/mouse). l) Representative histology showing the lung of IgG or IFNAR1 treated animals. m) Quantification of the mean linear intercept (MLI) from the lungs of RA and CS animals treated with IFNAR1 antibody or IgG control. Each dot represents the average of 3 fields of view from one section. n) Cytokine quantification of the lung homogenate. o) Quantification of mitochondrial polarization levels in the infiltrating neutrophils in RA and CS treated or not with anti-IFNAR1. Statistical analysis was performed by one-way ANOVA followed by Tukey’s multiple comparison. p) Representative image of lungs from RA or CS animals treated with IgG or anti-IFNAR1. Neutrophils are displayed in magenta (Ly6G), nucleus displayed as blue (DAPI) and NETs depicted in cyan (H3 Cit). q) Quantification of H3 Cit mean intensity normalized by the total lung area (RA IgG n=8, RA IFNAR1 n=8, CS IgG n=7, CS IFNAR1 n=7, animals from the first experiment batch). Data passed the Shapiro-Wilk test normality and statistical analysis was performed by One-way ANOVA followed by Tukey’s multiple comparison. Data variation is presented as ±SEM.

Therefore, we first used an acute Dextran Sulfate sodium (DSS)-colitis model (Fig. 5a). We found that mice treated with the IFNAR1 antibody displayed lower numbers of circulating neutrophils (Extended Data Fig. 5a). These cells displayed lower activation status, depicted by high CD62L and low CD63 levels (Extended Data Fig. 5b-c), and displayed lower mitochondrial potential (Extended Data Fig. 5d). Suggesting that blocking IFNAR1 may cause a short circuit to the IFN-α-IFNAR1 feedback loop in the circulating neutrophils.

In the colon, the IFNAR1 antibody treated animals showed lower weight to length colon ratio (Fig. 5b), improved (lower) histology score - based on ulceration, damage and leukocyte infiltration - (Fig. 5c), and lower crypt loss (Fig. 5d-e and Extended Data Fig. 5d). Anti-IFNAR1 treated animals also exhibited a better disease activity index (DAI) at earlier time points of the treatment (Extended Data Fig. 5f), less body weight loss (Extended Data Fig. 5g), less neutrophil infiltration (Fig. 5f-g) and NETosis (Fig. 5h-i). However, by the end of the DSS-colitis experiment, we did not observe changes in the DAI (Extended Data Fig. 5f). This may suggest that DSS-colitis, which is a model that directly compromises the epithelial integrity, may be too acute to observe significant improvements in DAI at later stages due to the damaged epithelium being the initiating insult. Collectively, our data suggested that the anti-IFNAR1 treatment decreased the number of circulating neutrophils and their activation in the DSS-colitis model.

Next, we employed a cigarette smoke (CS)-induced COPD mouse model to investigate the presence of IFNAR1+ neutrophils. We exposed mice to CS twice per day, 5 days per week for 2 weeks (Extended Data Fig. 5h). We observed that CS-treated mice displayed increased numbers of IFNAR1^+^ neutrophils in their lungs (Extended Data Fig. 5i). These neutrophils were not only active (IFNAR1^+^CD62L^-^) based on the CD62L shedding (Extended Data Fig. 5i), but also displayed higher mitochondrial polarization (Extended Data Fig. 5j). Supporting the existence of this population, found in the human scRNAseq analysis, in the CS model and the in vivo link between mitochondrial polarization and neutrophil activation.

We then sought to explore if the IFNAR1 blockade could inhibit neutrophilic inflammation and lung pathology in this model. Wildtype animals were treated with anti-IFNAR1 or IgG control monoclonal antibodies, and exposed to CS or room air (RA) (Fig. 5j). IFNAR1 treated animals displayed less neutrophils in their lungs (Fig. 5k, Extended Data Fig. 5k). We also observed decreased numbers of monocytes (Extended Data Fig. 5l), which may be related to the fact that neutrophils produce chemokines that attract other innate immune cells^40^. We observed no differences in the bone marrow or blood of these animals, suggesting that the increased numbers of infiltrating neutrophils may reflect higher migration, rather than increased granulopoiesis (Extended Data Fig. 5m-n). In addition, the IFNAR1 antibody-treated mice also displayed lower mean linear intercept (MLI) between their alveoli (Fig. 5l-m), and decreased levels of inflammatory cytokines in lung homogenate (Fig. 5n), including KC, TNFα, MCP-1, IL-6 and IL-12 (Fig. 5n, Extended Data Fig. 5o). Notably, we observed increased levels of IFN-α in the lung homogenate from CS mice, which was damped in the IFNAR1 treated animals (Extended Data Fig. 5p). Corroborating these findings, the infiltrated neutrophils had a lower mitochondrial polarization (Fig. 5o), and low activation status based on CD62L, CD63 and CXCR4 expression (Extended Data Fig. 5q-r). We also observed decreased NETosis in the smoked animals treated with anti-IFNAR1 (Fig. 5p-q). This data is in line with previous reports indicating that the *ifnr1*-/- mice have low MPO activity in their lungs^41^. Collectively, showing that anti-IFNAR-1 treatment can inhibit neutrophil metabolism and lung inflammation in a mouse model of COPD.

### Disrupting the IFNAR1 feedback loop mitigates neutrophil-mediated pathology in human organoid models of intestinal and pulmonary inflammation

Given the molecular and functional differences between murine and human neutrophils^38^, and that the monoclonal IFNAR1 treatment does not specifically target neutrophils, we sought to investigate the molecular cues driving tissue inflammation employing a human system. First, we validated that upon TLR9 agonist treatment, depleted *IFNAR1* neutrophils displayed lower degranulation and ROS, but phagocytosis remained unchanged (Fig. 6a). Similar results were found when we treated the neutrophils with an IFNAR1 monoclonal antibody in vitro (Fig. 6b). These findings were particularly interesting given that MyD88i-treated neutrophils displayed impaired phagocytosis (Fig. 6a); a function that is essential for maintaining the tissue homeostasis, while *IFNAR1* depleted neutrophils kept this function.

**Figure 6.**
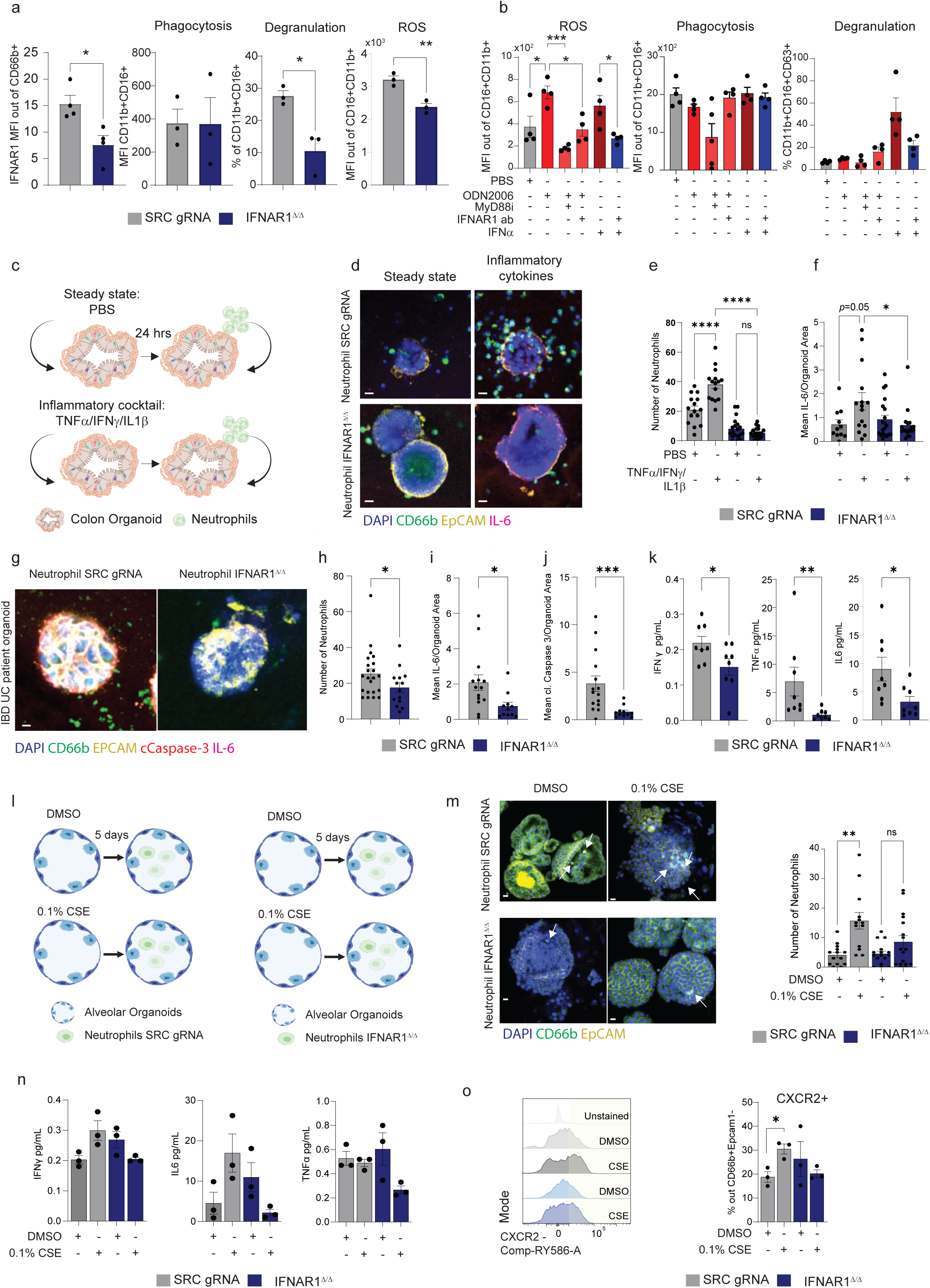
IFNAR1 depletion specific in neutrophils lead to impaired neutrophil inflammation and infiltration in intestine and lung organoids. a) Quantification of IFNAR1 signal upon depletion in fresh isolated neutrophils (left panel) and right panels shows the functional characterization under control (SRC, scramble gRNA, grey) and *IFNAR1* depleted fresh isolated neutrophils (navy). b) Validation of pathogenic neutrophil functions mediated by IFNAR1. MyD88i: MyD88 pan-inhibitor. The dots represent different donors. Data variation is presented as ±SEM. The “+” under the bar represents the addition of the compound. c) Graphic scheme showing the experimental design for the co-culture of neutrophils with colon organoids. Human organoids derived from healthy colon were cultured under steady state or exposed to an inflammatory cocktail to mimic an inflammatory milieu (Extended Data Fig. 6). After 24hrs organoids were retrieved from the domes and a new dome was made with neutrophils transfected with control guide RNA or *IFNAR1.* d) Representative imaging showing the neutrophil infiltration and IL-6 production in colon derived organoids and neutrophils co-culture. e) Quantification of neutrophils present within the colon organoid and f) IL-6 levels. Data generated from 8 neutrophils donors and 2 organoid donors. g) Representative imaging showing the neutrophil infiltration, cleaved caspase 3 (cCaspase-3) and IL-6 production in colon IBD derived organoids. h) Quantification of neutrophils present within the colon organoid, i) IL-6 levels and j) cleavage caspase 3. Data generated from 6 neutrophils donors and 2 organoid donors. Dots in the bars depicted a single organoid. From 2 independent colon donors and 8 neutrophil donors. k) Cytokine measurements from the supernatant of the IBD and neutrophils co-culture. l) Graphic scheme showing the experimental design for the co-culture of neutrophils with alveolar organoids (AO). Human organoids derived from healthy tissue were cultured under steady state (DMSO) or exposed to 0.1% cigarette smoke extract (CSE) for 5 days to mimic a smoked lung model (Extended Data Fig. 6). m) Representative imaging showing the neutrophil infiltration in alveolar organoids (left panel) and quantification of neutrophils present within the alveolar organoid (right panel). Dots in the bars depicted a single organoid from one AO donor and 3 neutrophil donors. n)Cytokine measurements from the supernatant of the AO and neutrophils co-culture. o) Left panel: Representative flow cytometry histogram showing the CXCR2 levels in CD66b+ (neutrophils) co-cultured with DMSO or CSE treated organoids. Right panel: Quantification of CXCR2 in neutrophils. Data from 3 neutrophil donors. Data variation is presented as ±SEM.

Next, we investigated if the *IFNAR1-*depleted neutrophils could dampen the inflammatory milieu characteristic of IBD and COPD. To this end, we developed 3D human organoid-based models for IBD and COPD, in which colon or alveolar organoids (AO) were co-cultured with human neutrophils, transfected with control sgRNA or sgRNA targeting *IFNAR1*.

For the IBD modeling, colon organoids were pre-treated with a cytokine cocktail (IFNγ, TNFα, and IL-1β) to mimic the inflammatory environment^38,39^ (Fig. 6d). This treatment increased MCP-1 secretion, epithelial permeability, and neutrophil migration (Extended Data Fig. 6a–d), confirming the establishment of an inflamed state. When co-cultured, *IFNAR1*-depleted neutrophils displayed reduced migratory capacity and IL-6 production compared with controls (Fig. 6e-f). Using patient-derived IBD organoids, we observed consistent findings: *IFNAR1*-deficient neutrophils showed reduced migration with fewer apoptotic events alongside lower IL-6, TNFα and IFNγ release (Fig. 6g–k). These results support a role for neutrophil-intrinsic IFNAR1 expression in sustaining intestinal inflammation.

To extend this to COPD, we generated a 3D alveolar model by exposing AO to cigarette smoke extract (CSE) for 5 days (Fig. 6l). CSE exposure enhanced neutrophil migration into the AO (Extended Data Fig. 6e-f), upregulated CXCR2 expression (Extended Data Fig. 6g-h), and increased alveolar cell apoptosis without affecting neutrophil viability (Extended Data Fig. 6i-j), thereby recapitulating COPD-associated pathology. We then employed this platform to test *IFNAR1* depleted neutrophils. We observed that *IFNAR1*-deficient neutrophils exhibited impaired migration (Fig. 6m; Extended Videos 3–6; Extended Data Fig. 6k), reduced IFNγ, IL-6, and TNFα release (Fig. 6n), and lower CXCR2 expression (Fig. 6o), and failed to promote alveolar cell death, in contrast to controls (Extended Data Fig. 6l). We also observed that neutrophils cultured with CSE-treated AO showed higher IFN-α levels, while the *IFNAR1* depleted neutrophils failed to produce IFN-α (Extended Data Fig. 6m-n), thus suggesting the presence of a positive feedback loop mediated by IFN-α that binds to IFNAR1 and may lead to the maintenance of inflammatory response in the tissue.

Collectively, our findings demonstrate that IFNAR1-expressing neutrophils contribute directly to IBD/COPD-related tissue damage in humans.

## Discussion

In this study, we uncovered shared neutrophil-driven inflammatory mechanisms between IBD and COPD by integrating scRNA-seq datasets and performing functional validation using an in vitro human neutrophil model. By employing the functional screen we identified a central role for mitochondrial metabolism, particularly regulated by *SLC25A37* (mitoferrin-1), which established an autocrine type I interferon (IFN-α) loop that amplifies neutrophil activation through IFNAR1 signaling. Finally, we confirmed this mechanism using human 3D organoids spiked with healthy or *IFNAR1* depleted neutrophils and provide insights into a therapeutic approach employing two different *in vivo* models that recapitulate key features of IBD and COPD.

Recent studies have identified distinct neutrophil subsets characterized by unique phenotypic markers and specialized functions^9,42^. Among the factors influencing neutrophil heterogeneity, a growing body of evidence suggests that neutrophil metabolism may be a key driver of this functional diversity^1,41,43^. Under homeostatic conditions, neutrophils primarily depend on glucose metabolism via glycolysis and the pentose phosphate pathway to meet their energy demands and produce NADPH^44–46^, which is essential for their antimicrobial effector functions^47^. However, in inflammatory diseases, this metabolic profile can shift. For instance, unstimulated neutrophils isolated from patients with IBD exhibit increased oxidative metabolism compared to healthy controls^48^. Interestingly, in CD patients, this enhanced oxidative activity negatively correlates with disease severity, suggesting a potential compensatory or regulatory mechanism that diminishes as inflammation progresses^49^.

Similarly, in COPD, intracellular NAD biosynthesis has been implicated in regulating neutrophil development, activation, and survival, potentially contributing to chronic inflammation^50^. Our data indicated that mitochondrial fusion and iron uptake are critical for maintaining elevated oxygen consumption and mitochondrial polarization. Taken together this data suggests that during chronic inflammation neutrophils increase the expression of *SLC25A37* (mitoferrin-1), a mitochondrial iron transporter, in order to promote mitochondrial fusion and oxidative phosphorylation (OxPhos).

Mitochondrial dynamics are increasingly recognized as key modulators of innate immune cell function, particularly in response to infection stress. Mitochondrial fusion, a process orchestrated by mitofusin-1 (*MFN1*), mitofusin-2 (*MFN2*), and optic atrophy 1 (*OPA1*)^51^, has been implicated in the regulation of NET formation. OPA1-dependent ATP production is essential for NETosis and is critical for host defense against *Pseudomonas aeruginosa*^52^. Beyond bacterial infections, mitochondrial function also plays a pivotal role in antiviral responses^53–55^. Mitochondrial antiviral-signaling protein (MAVS) is central to viral sensing^56,57^, and recent evidence has demonstrated that mitochondrial fusion is regulated by the SIRT3–OPA1 axis, which is essential for defense against human cytomegalovirus (HCMV)^58^.

In this study, we demonstrate that neutrophils are capable of sensing NET-derived products through TLR9, a process that elicits mitochondrial OxPhos. Specifically, mitoferrin-1-mediated mitochondrial fusion enhances OxPhos, promoting the production of type I interferons via TLR9 signaling. Accordingly, in tuberculosis it was reported that type I IFN-mediated NET release is associated with granuloma caseation^59^ and are capable of activating pDCs through the release of NETs, driving a chronic pDC activation facilitating the onset of autoimmunity in systematic lupus erythematosus (SLE)^60^. Indicating that IFN-α may trigger neutrophils towards a “pathogenic” phenotype that can promote inflammation in the tissue. Consistently, it has been reported that CS can promote neutrophil-CXCL8 production in a TLR9 activation-dependent manner^61^ and neutrophils isolated from SLE patients displayed a proinflammatory phenotype that synthesizes type I IFNs^24,62^. These findings identify a novel mechanism by which sterile inflammation may be driven by mitochondria-fueled NET sensing.

Type I IFNs signal through specific heterodimeric receptors: IFN-ɑ/-□ (IFNAR) that are comprised of IFNAR1 and IFNAR2. Upon activation the IFNAR1 dimerizes and triggers the tyrosine kinase 2 (TYK2) leading to the phosphorylation of STAT1 and STAT2. These are the driving transcriptional factors, which will activate the nuclear complex termed IFN-stimulated gene (ISG)^63^. Interestingly, TYK2 impairment by the rs34536443 and rs12720356 SNPs are reported to have protective effects against IBD^63,64^. However, in contrast to our IFNAR1 antibody treatment, global *Ifnar1* mice are more susceptible to DSS-colitis, however, during the recovery phase of DSS colitis, global loss of IFN-I signaling favors tissue repair due to decreased epithelial cell apoptosis and lower monocyte migration^63^. This suggested that IFNa-IFNAR1 axis dynamics may differ depending on acute versus chronic inflammation.

Collectively, our findings highlight a mitochondrial–NET–TLR9–type I IFN axis that contributes to mucosal inflammation. This pathway may represent a previously unappreciated mechanism by which sterile triggers initiate chronic inflammatory responses, and offers a potential therapeutic target in diseases characterized by excessive NET formation and type I IFN activity such as airway inflammatory conditions.

## Methods

### Animals

Animal studies were approved by the relevant local regulatory authorities (BS Kantonales V *Sampling of mice:* terminal blood collection, lung perfusion followed by lung, spleen and bone collection Anesthesia: terminal anesthesia, 150 mg/kg body weight pentobarbital diluted in 0.9% NaCl physiol. I.p.

#### Lung collection and homogenate

Veterinäramt BS, cantonal approval number #3065 and #3090; national number #35788), and all experimental protocols adhered to both federal and cantonal regulations. The experiments were carried out in AAALAC-accredited facilities at F. Hoffmann-La Roche in Basel. Mice were housed under specific-pathogen-free conditions with continuous health surveillance throughout the studies.

#### CS Model

A total of 30 female C57BL/6J mice from CRL-F of the age of 9w were included in the study. Having 7 animals for the CS group and 8 animals for the RA group housed 3-4 animals per cage.

#### Smoking challenge

Mice were exposed to whole body mainstream CS generated from 1R6F research cigarettes (29.1 mg tar/1.896 mg nicotine) using the following standard parameters (ISO 1991): one 60 ml puff of 2 seconds duration followed by 58 seconds of fresh air. The cigarette smoke is directed in the exposure chamber (5 liters volume). Flowrate is constant at 0.833 L/min.

#### Animal treatment

IFNAR1 inhibiting antibodies (P1AQ1038) or isotype control (P1AQ1040) were dosed twice per week via 200uL i.p. injection. Therefore, the concentration of the solution was 2.5mg/mL. Mice were monitored and weighed every day during the smoking exposure. Mice were assessed for difficulty in breathing, activity, posture, general behavior and body weight loss in the appropriate score sheet.

Lungs were carefully isolated without the trachea and esophagus and split as follows:

a. Left lobe of lung for flow cytometry. Samples were kept on ice on a 24well plate in PBS + EDTA.
b. Superior, middle and inferior lobe for homogenization for cytokine and chemokine assay.
c. Post-caval lobe was isolated into histocasette and then kept in 10% Formalin for 24h before transferring to 4°C PBS

#### Lung homogenization

Lungs for *b* were weighted and snapped frozen in a 7mL Precellys tube. To each sample 100 µL of lysis buffer (200 mM NaCl, 5 mM EDTA, 10 mM Tris, 10% glycerin and Roche cOmplere protease inhibitors) was added per 10 mg tissue. The samples were then homogenized in the Precelly tissue homogenizer by 5500 rpm, 3 x 20 sec and 20 sec pause. After, the homogenate was transferred to a 2 mL eppendorf tube and centrifugated for 15 min at 1500 x g at 4°C. The supernatant was removed and transferred to a new 2 mL Eppendorf tube and centrifugated at 4°C at 21000 x g for 10 min. The supernatant was transferred on a 96-well plate and stored until the Meso Scale Discovery assay was performed (MSD).

#### MSD U-R-Plex

To evaluate the inflammatory cytokines IL-12, IL-6, IFN-γ, Rantes and TNF-α we used the MSD U-plex Custom biomarker (Cat. No. K15069M-1; Lot: 504404).

#### Bone marrow collection for smoking model

For flow cytometry, mouse bone marrow cells were isolated from pooled femur and tibia by flushing them with cold phosphate-buffered saline (PBS) containing 2% FCS. The fluxed material was filtered in a 40 mm filter and transferred in a 50 mL conical tube. The samples were centrifugated for 5 min at 350 x g at 4°C. The supernatants were removed and 5 mL of ACK Lysing Buffer (Thermo Fisher Scientific) was added. The cells were then resuspended in FBS-BD-FACS staining buffer and washed twice by 5 min at 350 x g at 4°C centrifugation. About 1 million cells were incubated with an antibody cocktail containing LyG6 (clone 1A8, Biolegend), Ly6C (clone HK1.4, Biolegend), CD11b (clone M1/70, Biolegend), CD62L (clone MEL14, Biolegend), IFNAR1 (clone MAR1-5A3, Biolegend), CD63 (clone NVG2, Biolegend), CD45 (clone S18009F, Biolgened), Lineage (F4/80 clone BM8, CD3 clone 17A2, B220 clone RA3-6B2) and viability dye. Samples were acquired using an Aurora Cytek cytometer.

#### Dextran sulfate sodium (DSS) colitis model

A total of 24 female C57BL/6J mice from CRL-F of the age of 9w were included in the study. The animals were separated into 3 groups (8-7 animals per group). The DSS challenge was performed by adding 2.00% of DSS (SKU 0216011090, MP Biomedicals) to the mice drinking water.

#### DSS-Colitis Treatment

IFNAR1 inhibiting antibodies (InVivoMab, Bio X Cell, P1AQ1038) or isotype control (P1AQ1040) were dosed twice per week via 100uL i.p. Injection (500µg/100µL).

#### Daily scoring (Day 4-12

Animals were placed into an empty cage to assess fecal samples, during this time the body weight was taken and the animals were scored. To assess severity of colitis the Disease Activity Index (DAI) was calculated as a sum of the scores of body weight loss, stool consistency and blood in stool.

#### Tissue collection

For the blood collection after euthanisia we retrieved about 500-700µl from the right ventricle. The blood was transferred to an EDTA tube. The syringes were also previously coated with PBS-EDTA.

For the colon we dissected the gut and separated the small intestine from the colon. Next we removed fecal pellets and selected one centimeter of the mid to distal section for histology and kept them in 4%-PFA overnight, then transferred to PBS for at least 24h until further embedding in paraffin.

#### AI-Based Histological Image Analysis for the colon sections

Transverse sections of the distal colon were stained with H&E as previously described. Images were acquired at 40x magnification NA missing, producing one digital image per mouse, with 1-4 tissue sections per image.

Digital image analysis was performed using the commercial software Visiopharm (Version 2024.07.1.16912 x64). The workflow was developed to reproduce manual scoring criteria described by Smith et al. ^66^ and Dieleman et al. ^66,67^, focusing on histological features relevant to colitis, specifically ulceration and crypt damage.

The analysis pipeline was internally validated by comparison to manual scoring across multiple dextran sulfate sodium (DSS)-induced acute colitis studies. Each step of the pipeline was optimized through an iterative process: representative fields of view were manually annotated, artificial neural networks were trained, and performance was assessed on randomly selected images. A step was considered validated if visual confirmation of results reached ≥90% accuracy. Once validated, the step was applied to at least 50 images from various studies and disease states. If performance did not meet criteria, further manual annotations were added, networks were retrained, and post-processing steps were adjusted. This iterative process was repeated until consistent alignment with ground truth was achieved.

#### Detailed image analysis workflow

a. *Tissue segmentation:* Whole transverse tissue sections were detected using a DeepLabv3+ neural network at 1x digital magnification. The model was trained on two classes (background, tissue) with 76,891 iterations and a final loss function value of 0.001. Detected tissues were enumerated to determine tissue count per image. All downstream analyses were performed on individual tissue sections, and final per-mouse values were calculated as the mean of all tissue sections.
b. *Layer detection:* Within each tissue region of interest (ROI), anatomical layers were identified using a DeepLabv3+ network trained with 112,000 iterations and a final loss of 0.017. Images were analyzed at 4x virtual magnification with four classes: background, lumen, mucosa, and gut-associated lymphoid tissue (GALT). GALT regions were excluded from all further analyses.
c. *Crypt detection and scoring:* Within the mucosa, colon crypts were identified using a DeepLabv3+ network trained with 78,840 iterations and a loss value of 0.022. Images were processed at 10x virtual magnification using three classes: crypt area, crypt loss, and GALT. GALT regions were again excluded to ensure specificity. This step allowed the quantification of intact versus damaged crypt areas, enabling calculation of the percentage of crypt loss within the mucosa.
d. *Ulceration scoring:* Ulceration was quantified by measuring the total interface length between the lumen and mucosa and comparing it to the length of interface where crypt loss directly abutted the lumen—indicating a loss of epithelial barrier and presence of ulceration. The ratio of ulcerated interface to the total lumen-mucosa interface was used to assign scores:

- <1%: Score 0 (no relevant ulceration)
- 1–10%: Score 1 (light ulceration)
- 10–50%: Score 2 (moderate ulceration)
- 50%: Score 3 (severe ulceration)

### Immunofluorescence staining

Tissue immunofluorescence staining was performed using the *Ventana platform*. Initially, the slides were pre-warmed to 60 °C and incubated for 8 min (baking). This was followed by deparaffinization, which consisted of three cycles of warming at 60 °C for 8 minutes each.

Subsequently, the Cell Conditioning 1 (CC1) protocol was initiated, during which slides were heated to 95 °C and incubated for 4 minutes. A gradual temperature ramp was applied, increasing the incubation time by 8-minute increments until a total of 88 minutes was reached.

Following CC1, the *DISCOVERY inhibitor* protocol was applied. Primary antibodies were then manually added: Ly6G (CST, #87048s, 1:50 dilution) and H3Cit (Abcam, #ab5103, 1:50 dilution). The slides were incubated with the primary antibodies at 37 °C for 60 minutes.

Next, the [OMap anti-Rabbit HRP protocol] (One Drop) was selected and incubated for 16 min. This was followed by the addition of DCC H₂O₂ (One Drop), with a 20-min incubation.

The Dual Sequence (DS) Cell Conditioning 2 (CC2) protocol was then carried out. Slides were heated to 100 °C and incubated for 4 min, followed by three 8-min increments for a total incubation of 24 min.

Secondary antibodies were then applied. The slides were gradually warmed to 37 °C from low temperature and incubated for 60 min. The DS Multimer HRP method was used next, followed by the [OMap anti-Rabbit HRP] protocol (One Drop), incubated for 16 min, during which the DS Cy5 detection system was employed. A Cy5 H₂O₂ reagent (One Drop) was then applied and incubated for 20 min. Finally, DAPI was used as a nuclear counterstain.

### Cell typist analysis

For the harmonization of the IBD and COPD scRNAseq from published available dataset, the custom model option was used from CellTypist ^68^. To this end we trained a “neutrophil” module based on the neutrophil profile described in ^42^, we also used the pre-made module “immune cells” version 1 model.

### In vitro neutrophil differentiation

*Progenitor cell expansion:* CD34+cells (Lonza) from adult or cord blood were thawed in X-VIVO. The samples were centrifuged at 300 g for 10 min to remove the freezing solution and resuspended in Progenitor cell expansion media (PEM) containing X-Vivo media (Lonza), 20 % BIT Serum (Stem cell), SCF (100 ng/mL), IL-3 (50 ng/mL), FTL3L (50 ng/mL), IL-6 (50 ng/mL), G-CSF (100 ng/mL), TPO (10 ng/mL), insulin-Transferin-Selenium (1x) and DNAseI (1%). The progenitor cells were cultured in 96-U bottom plates in 100 uL PEM.

#### Myeloid polarization

After 3 days in culture the wells containing the progenitor cells were combined in a 50 mL falcon tube and centrifuge at 300 g for 10 minutes (break off). The pellet was resuspended in the Myeloid polarization media (MPM), containing X-VIVO, 20 % BIT, SCF (100 ng/mL), G-CSF (10 ng/mL) and DNAseI (1%). The cells were plated into 96-well plates (10.000 cells/well) with 100 uL MPM.

#### Neutrophil polarization

After 4 days in culture with MPM the wells were combined each well in a 15 mL falcon tube and centrifuged at 300 g for 10 min (break off). The pellet was resuspended in Neutrophil media having G-CSF (100 ng/mL), DNAseI (1%), Glutamine (0.1%),Essential Amino Acid (0.1%) and Rosiglitazone (0.1 ng/mL).

All the cytokines were purchased from Peprotech Proteins| Thermo Fisher Scientifics.

The culture was expanded to 6 well-plates containing 2mL of neutrophil media (0.8 x 10^6^ cells). After 15-17 days we used the stemcell negative neutrophil selection following manufacturing recommendations (StemCell technology) and evaluated it based on the cell morphology and flow cytometry (CD66b, CD11b and CD34).

### CRISPR-Cas9 mutagenesis

Target selection for CRISPR-Cas9–mediated mutagenesis was performed based on Human GecKO library guides ^69^. The selected sgRNA having a GC content <55%, mismatches (MM) self-complementarity = 0, MM0 ≤ 1, MM1 ≤ 1, MM2 ≤ 1, MM3 ≤ 1 were selected. The sgRNA templates were generated using the protocol described by Gagnon *et al.* ^70^ using the mMESSAGE mMACHINETM SP6 transcription kit (Invitrogen, AM1340). Editing efficiency was further validated by deep sequencing of PCR amplicons.

For the transfection of the guides ∼ 100.000 neutrophils were resuspended in P3 Primary solution (Lonza) containing the sgRNA (20 μM) and Cas9 Protein (TrueCut v2.0, 10 μM) and placed in 16 wells-Stripes cuvettes from Lonza. The cells were then exposed to the ZD100 program and after transfection 100 μL of X-VIVO-20% BIT was added to the wells. The cells were kept untouched for 5 min at room temperature.

Amplification was performed using Phusion High-Fidelity PCR Master Mix with HF Buffer (New England Biolabs, catalog no. M0530) under the following conditions: 98°C for 3 min, 98°C for 10 s, time for annealing for 10 s, and 72°C for 10 s × 35 cycles, and then 72°C for 5 min. Then, PCR amplicons were purified on PCR purification kit columns before sequencing and sent to MicroSyth for the Sanger sequencing. The sequencing results were analysed using the *Tide* ^71^ software.

### NETosis imaging

After neutrophil differentiation and isolation using the Stem Cell negative selection we prepared a neutrophil stock of 200.000 cells/mL in culture media (X-Vivo Lonza, 10%-Plus Serum alternative BIT from StemCell). The cells were then cultured in 0.1%-Gelatin precoatted 96-well glass bottom plates in the concentration of 40.000 cells per well. The cells were allowed to settle at room temperature for 15 minutes. After, the plate was centrifugated at 70 g for 3 min (no break) and 40 uL of Sytotox Green Dye Solution added to a final concentration of 250 nM, from my stock 1: 1000 Media (X-Vivo Lonza). As positive control we added 10 uL of fMLP (Sigma, 10 uM), or LPS (Sigma). For negative controls we used 1% DNAse 1 (Roche) or the PADI4 inhibitor (Chloramine, 10 uM).

For the live imaging the plate was incubated in the Incucyte (Sartorius) and images were taken for 6 hrs with an interval of 20 min/30 min.

For the antibody staining 4 hrs after challenge the supernatant was carefully removed and 100 uL of the BD fix buffer was added. The plate was kept in the fridge overnight. Then, the supernatant was removed and the plate washed with PBS by spinning the plate at 100g for 3 min. After the wash, we added a FBS-FACS BD Staining buffer containing anti-Histone H3 (citrulline R2 + R8 + R17) (Abcam, RM 1001) coupled with Zenon (ThermoFisher, Z25402), CD63 (FITC, BD AB_393869) and DAPI. The plate was incubated at room temperature for 2 hrs. After, the plate was washed twice with PBS and images were taken using the Leica LAS X confocal microscope.

The functional characterization after the CRISPR-Cas9 perturbation was conducted 12 hrs after the transfection.

### Seahorse Real-Time Cell Metabolic Analysis

For the real-time cell metabolic assay the Seahorse XFp Cell Mito Stress Test Kit was used following the manufacturer recommendations. In brief, fresh isolated neutrophils or transfected neutrophils were plated as 60.000 cells/0.1% coated Gelatin wells. After plating the cells were stimulated with the TLR agonists (indicated in the Figures) or with PMA as positive control and immediately placed into the prepared Seahorse XF Pro Analyzer. The total of 3-5 cycles were settled for the baseline.

### Simple Western Automated Western Blot System (WES)

Twelve hours after transfection the human neutrophils were collected into 50 mL conical tubes and spinned at 350g for 5 min (no break). After, the pellet was resuspended in RIPA Lysis and extraction buffer (ThermoFisher) and total protein concentration evaluated using the Pierce^TM^ Bicinchoninic Acid (BCA) protein assay (ThermoFisher). Then, WES assay was conducted following the product recommendation. The total of 10 μg protein was loaded into the WES capillaries. The SLC25A37 (1:10 dilution, Invitrogen ref:PA5-26720) and Vinculin (1:200 dilution, Cell Signaling ref: 13901) were used.

### Flow cytometry and confocal microscopy

Fluorochrome-conjugated monoclonal antibodies were purchased from eBioscience, BD Pharmingen, or Biolegend and staining performed as previously described ^72^. Fresh isolated neutrophils, transfected primary neutrophils or in vitro derived neutrophils were quantified by direct staining with MitoTracker (Red CMX and Green^TM^, ThermoFisher), CMxROX (deep red, ThermoFisher), and mito probe JC-1 staining was performed according to the manufacturer’s instructions. Intracellular cytokines production in the fresh isolated neutrophils were quantified 4 hrs after stimulation with TLRs and GolgiStop (BD). The cells were fixed and permeabilized using the transcription factor staining buffer set (eBioscience). For the phagocytosis experiments fresh isolated neutrophils or in vitro derived neutrophils were stimulated with fMLP (Sigma), LPS (Sigma) or ODN2006 (Invitrogen) and incubated with the S.Aureus-pH-Rhodo or E.Coli-pH-Rhodo (ThermoFisher) for 4 hrs. Cells were acquired on Fortressa flow cytometers (BD Biosciences) or Aurora Cytek and analyzed using FlowJo (TreeStar) software. Cells were imaged live on glass bottom dishes coated with 0.1% Gelatin using a LAS X Leica confocal scanning microscope (Leica) or with the Incucytes (Sartorious) using the non-adherent protocol and a 20x Nikon objective. Cells were kept in a humidified incubation chamber at 37°C with 5% CO2 during image collection. Images were deconvolved and analyzed using ImageJ (NIH).

### Proteomics

One million fresh isolated neutrophils, in vitro derived neutrophils, HL-60 and HL-60 stimulated with Phorbol myristate acetate (PMA; 1μg/mL) were resuspended in 50 μL of LYSE-iST buffer (PreOmics kit, ref: P.O.00027) containing 0.5 μL TCEP 5mM final (Bond-Breaker, ThermoFisher ref: 77720). The cells were incubated in a heating block at 95°C, 1000 rpm for 10 min. After, the samples were sheared in the Bioruptor (10 cycles, 30 sec ON/OFF). The total protein concentration was quantified with the BCA protein assay (ThermoFisher). Digestion and purification were conducted following the PreOmics kit instructions. Next we performed by the peptide assay following the Peptide Assay (ThermoFisher; ref: 23275) instructions. The samples were resuspended in LC-MS resuspension buffer (2% ACN-0.5% FA).

For the MS measurements 7 μL from each 0.2 μg/μL sample were transferred to a 96 well plate. For the MS measurements the Trap column DNV pepMap (Neo ref:DNV75150PN) and the 25cm Aurora Series emitter column; (25cm x 75µm ID, 1.7µm C18) (Ion Optiks; # AUR3-25075C18; Lot#IO2575028066) were used. The samples were acquired in the Vanquish Neo LC system and an Orbitrap Tribrids Ascend MS System. The data were analyzed using the Spectronaut software. Samples were injected randomized and ∼ 4.881 proteins identified.

### RNAseq bulk

RNA was extracted using the MiniRNA Isolation Kit (Thermo Fisher Scientific). The quality between was evaluated using the Fragment Analyzer. RNA that passed the quality control (QC) was then used to generate libraries using the NEBNext Library Ultra Low Input Library Preparation. The manufacturer’s recommendations were followed, and the libraries were sequenced on an Illumina NovaSeq6000 sequencer. All sequencing data were performed in three biological replicates, having a total depth of 15 million reads per reads. The reads were conducted at 2 × 150 base pair (bp) of length.

### Bulk RNA-seq bioinformatic analysis

Reads for mouse bulk RNA-seq datasets were mapped following the default Biokit 3.9 (pRED workflow). Briefly, FASTA files were aligned against the human genome version GRCh38. Reads were counted with featureCounts. Differential expression analysis was performed with DESeq2. Multi-FASTA QC statistics–indicated data were of high quality, and sequencing depth was sufficient to test for differential expression between conditions. Differentially expressed genes were called with a FDR threshold of 0.05. Correlations between datasets and graphical output were conducted in an R environment and CellTypist.

### Enzyme-linked immunosorbent assay (ELISA)

The human Citrullinated histone H3 (H3Cit) was quantified from the supernatant of the in vitro derived neutrophils stimulated for 12 hrs using the H3Cit ELISA kit (Cayman Chemicals, ref: 501620).

Secreted IFN-α from 4 hrs stimulated fresh isolated neutrophils or the mouse tissue homogenate was quantified with the Human IFN-α ELISA Kit (ThermoFisher, ref:BMS6027) or the Mouse IFN-α ELISA kit (ThermoFisher, ref:BMS6027).

#### Tissue source

Human tissues and associated clinical information were obtained from patients undergoing tumor resection at the Humanitas Research Hospital, Milan, Italy. Samples were sourced through a collaborative framework adhering to legal and ethical regulations. The project was reviewed and approved by Comitato Etico Territoriale Lombardia 5 (Ethical approval 3631). Additional tissue samples and annotated data were obtained, and experimental procedures were performed within the framework of the non-profit foundation HTCR, which includes written informed consent from all donors and has been approved by the ethics commission of the Faculty of Medicine in the LMU (number 025-12) and the Bavarian State Medical Association (number 11142)

### Colon Organoids

#### Human sample preparation, crypt isolation and organoid culturing

Intestinal tumour-free regions were used for crypt isolation. First, the muscularis, serosa and fat from the basal side was removed with forceps and scissors. Mucus from the luminal side and blood vessels from the apical side were removed with a scalpel. The processed mucosal tissue was then washed five times with PBS supplemented with penicillin-streptomycin (Thermo, #15070063), gentamicin (Thermo, #15750060), amphotericin (Thermo, J67049-AD). For the isolation of crypts, a 3 cm^2^ piece was scrapped with a cover slip to remove the villi. The remaining tissue was then incubated in cold PBS supplemented with 10 mM EDTA for 30 min on a shaker. The crypts were afterwards scraped off with a cover slip and collected in organoid media (Advanced DMEM/F12 Gibco #12634028, HEPES Gibco #15630080 10 mM, Glutamax Gibco #35050061 1%, Wnt surrogate ImmunoPrecise Antibodies #N001-0.5mg 0.15 nM, Noggin-CM ImmunoPrecise Antibodies N002 1:50, Rspo3-CM ImmunoPrecise Antibodies #R001-500 ml 1:50, hIGF-1 BioLegend #590908 100 ng/ml, hFGF2/FGF-basic R&D #3718-FB 50 ng/ml, Gastrin I MedChem Express #HY-P1097/CS-0027717 10 nM, A83-01 (ALK4/5/7 inhibitor) R&D #2939 0.5 µM, hEGF PeproTech #AF-100-15-1MG 50 ng/ml, B27 min VitA Thermo #12587010 1:50, N-acetylcysteine Merck/Sigma #A9165-5G 1mM, Primocin Invivogen #ant-pm1 50 µg/ml). Crypts were centrifuged, the pellet was resuspended in matrigel (Corning, #356231) and after polymerization (15 min, 37°C) cultured in organoid media with 10 uM Y-27632 (STEMCELL Technologies, 72305). The organoids were passaged every week. Therefore, media was removed and after PBS wash, Gentle Cell Dissociation reagent (STEMCELL Technologies, #07174) was added for 2 min RT. The domes were collected in falcon tubes, further incubated for 15 min at RT and centrifuged. The pellet was resuspended in organoid-wash media (Advanced DMEM/F12 Gibco #12634028, HEPES Gibco #15630080 10 mM, Glutamax Gibco #35050061 1%, Primocin Invivogen #ant-pm1 50 µg/ml) and multiple times resuspended to break the organoids. This process was repeated before the pellet was resuspended in matrigel. After passage three, a mycoplasma test was performed (Lonza, #LT07-710) before the organoids were used for experiments.

For experimental preparation at day 3-4 after passing, the media was removed and after washing with PBS, the organoids were treated with cell recovery solution (Corning, CLS354253) and incubated for 40 min at 4°C. The organoids were transferred into a falcon tube, well resuspended and centrifuged.

#### IBD derived organoids establishment and expansion

Mucosal tissues were separated from the muscularis, serosa and fat using forceps and scissors. The processed mucosal tissue was then washed five times with PBS supplemented with penicillin-streptomycin (Thermo, #15070063), gentamicin (Thermo, #15750060), amphotericin (Thermo, J67049-AD). Tissue was minced into small pieces and subsequently enzymatically digested in 1 ml of advanced DMEM/F-12(GIBCO, #12634028) medium supplemented with 20 U/ml and 20 μg/ml of P/S (ThermoFisher Scientific #15140122), Primocin 0.1 mg/ml (InvivoGen, #ant-pm-1), Hepes 10 mM (GIBCO, #15630056), and 2nM Glutamax (GIBCO, #35050061), 2.5 mg/ml collagenase IV (Worthington, #LS004189), 0.1 mg/ml DNase IV (Sigma, #D5025), 20 μg/ml hyaluronidase V (Sigma, #H6254), 1% BSA (Sigma, #A3059), and 10 μM LY27632 (Abmole Bioscience, #M1817) for 1 h at 37°C under slow rotation and vigorous pipetting every 15 min.The tissue lysate was filtered through a 100 μM cell strainer and centrifuged at 300 g for 10 min. The cell pellet was suspended in PBS and cells were counted using trypan blue (GIBCO, #15250061), Countess™ Cell Counting Chamber Slides (Invitrogen, #C10228) in Countess™ II FL Automated Cell Counter (Invitrogen, #AMQAF1000). Cells were resuspend in growth factor reduced Matrigel (Corning, #356231) at a concentration of 8000 cells/ul and seeded as 20 ul drops in a tissue-culture dish.After polymerization of the Matrigel, IntestiCult(# 06010) medium was supplemented with 10uM Rock Inhibitor. Organoid medium was changed every 3 days and, when needed, organoids were passed after dissociation with Gentle Cell Dissociation reagent as previously described for Colon Organoids.

#### Coculture with neutrophils

Before co-culture the human colon organoids were gradually exposed to IntestiCult™ Intestinal Organoid Culture Medium (StemCell Technologies, #06010) until fully cultured using this medium. Healthy and patient derived-IBD, were maintained in IntestiCult™ Intestinal Organoid Culture Medium (StemCell Technologies, #06010) from thawing. After expansion, organoids were fragmented by mechanical shearing through repeated pipetting and pelleted by centrifugation at 400 x g at 4°C. The supernatant was removed and organoid fragments were resuspended in an extracellular matrix mixture composed of type I collagen (Ibidi, #5021) and Matrigel (Corning, #356231) at a 1:4 (v/v) ratio. Neutrophils from healthy donor peripheral blood were isolated by negative magnetic selection (StemCell Technologies, #100-0404) and immediately combined with organoid fragments in the matrix, targeting ∼30% neutrophils by input cell number per dome, reflecting proportions observed in inflammatory bowel disease^73^. The cell–matrix suspension was dispensed as 10-µL domes into glass-bottom 96-well plates and allowed to gel at 37 °C according to matrix manufacturer recommendations. Domes were overlaid with prewarmed IntestiCult media and maintained under standard culture conditions until downstream assays.

After 24 hrs, the medium was removed from wells and 4% Fixation solution (BD Cytofix, #554655) was added. The plate was then incubated at 4°C for 2 hrs. Afterwards, the fixation buffer was removed and the samples washed twice with stain buffer (FBS) (BD Biosciences, #554556). Then, organoids were blocked using 0.5% BSA in DPBS for 30min at room temperature. Supernatant was then removed and samples were resuspended in stain buffer supplemented with the primary antibodies CD66b (1:200, FITC, Biolegend, clone: G10F5, #984102), IL-6 (1:100, BD Pharmigen, APC, clone: MQ2-13A5, #561441) and EpCAM/CD326 (1:100, Alexa-700, Biolegend,clone:9C4, #324244) and incubated in staining buffer solution overnight at room temperature. On the next day, organoids were washed once in DPBS buffer and the plate was imaged in a LAS X Leica confocal scanning microscope (Leica) and a 40x Nikon air-objective. Images were deconvolved and analyzed using ImageJ (NIH)

### Alveolar Organoids

#### Culture expansion

Adult human pulmonary alveolar epithelial cells (ScienCell, #3240) were cultured within matrigel (Corning, #356231) domes, in which 24-well plates were used and one dome was seeded per well. Passaging of organoids was done every 10 days using the Animal Component-Free Cell Dissociation Kit (ACF, StemCell Technologies, #05426) to dissociate the organoids into single cells, as recommended by StemCell Technologies. Cells were centrifuged at 400 x g for 5 min at 4°C, and replated at a concentration of 1.6 x 10^5^ cells/mL. After 30 minutes at 37°C, PneumaCult Alveolar Organoid Expansion (AvOE) seeding medium (StemCell Technologies, #100-0847) was added to the domes, and changed every 3-4 days for normal AvOE medium.

#### Cigarette smoke extract (CSE) treatment and co-culture with neutrophils

On day 10, the alveolar organoids were treated with DMSO (Merck, #D138401) or CSE (Fraunhofer Institute for Toxicology and Experimental Medicine ITEM, generated from 1R6F cigarettes) 0.1% for 5 days in PneumaCult AvOE medium. On day 15, organoids were dissociated from the matrigel into fragments. In brief, gentle cell dissociation regent (GCDR, StemCell Technologies, #100-0485) was added for 2 min at room temperature. Then, domes were transferred to a 15 mL 1% BSA-coated falcon and incubated for 10 min at room temperature. After incubation, samples were centrifuged at 400 x g for 5 min at 4°C. The supernatant was removed and 10 mL of washing buffer (DMEM/F12+glutamax; Gibco, 30% BSA; Merck and HEPES; Gibco) was added. The organoids were then disrupted by mechanical forces through pipetting. Organoids were centrifuged as before, supernatant was removed and organoids were resuspended in matrigel in a 1:3 dilution and mixed with healthy neutrophils isolated from blood using the negative magnetic beads enrichment (StemCell Technologies, #100-0404). The proportion used was approximately 5% neutrophil to dome, which corresponds to 600K neutrophils for 12 domes. This proportion was chosen based on the ratio found in COPD patients^74^. The cell mix was plated into 25 µL-domes of matrigel into 24-well plates.

After 30 minutes at 37°C, 50% of PneumaCult AvOE seeding medium and 50% neutrophil medium supplemented with DMSO/CSE was added to the domes. The plate was then incubated at 37°C with 5% CO2 for approximately 24 hrs.

For flow cytometry, samples were first dissociated using the ACF kit, as previously described for passaging. Then, cells were centrifuged at 300 x g at 4°C for 5 min, washed in cold stain buffer (FBS)(BD Biosciences, #554556), and centrifuged as before. Then, the Zombie violet viability dye (Biolegend, #423113) and Human BD Fc Block (BD Biosciences, #564220) were added and the cells were incubated at room temperature for 15 min protected from light. Followed by washing in a cold stain buffer and centrifugation. Samples were then transferred to a 96-well U-bottom plate and labelled with primary coupled antibodies, EpCAM/CD326 (1:100, Alexa-700, Biolegend,clone:9C4, #324244), CD66b (1:200, FITC, Biolegend, clone: G10F5, #984102), CD62L (1:100, BUV615, BD, clone: SK11, #751364) and CXCR2/CD182 (1:100, RealYellow586, BD, clone:6C6, #753253) were added. The plate was then incubated at 4°C for 40 min. Afterwards, wells were washed with the stain buffer, centrifuged as before and the pellet was resuspended in a fresh stain buffer. Samples were filtered before acquisition using Aurora Cytek and analyzed using FlowJo software.

For immunofluorescence, the organoid-neutrophil mix was recovered from matrigel using cell recovery solution (Corning, #354253). Samples were incubated on ice for 40 min, and then centrifuged at 300 x g at 4°C for 5 min. Supernatant was removed and samples were fixed with 4% paraformaldehyde (PFA, MP Biomedicals, #0219998380) for 45 min at room temperature. Samples were centrifuged at 200 x g at room temperature for 5 min and washed with IF Buffer: 0.1% BSA (Sigma-Aldrich, #A9647), 0.2% Triton X-100 (Sigma-Aldrich, #T8787) and 0.05% Tween20 (Sigma-Aldrich, #P9416) in DPBS. Then, organoids were centrifuged and blocked using 5% donkey serum (Sigma-Aldrich, #D9663) and 1% Triton X-100 in DPBS for 1h at room temperature. Samples were centrifuged and primary antibodies mature surfactant protein C (SP-C, 1:500, SevenHills Bioreagents, #WRAB-76694), CD66b (1:200, FITC, Biolegend, clone: G10F5, #984102), CXCR2/CD182 (1:100, PE/Dazzle594, Biolegend, clone: 5E8, #320722) or IFN-ɑ (BD Pharmigen, PE, clone: 7N4-1, #560097) were incubated in a blocking solution overnight on a rocker at room temperature. On the next day, organoids were washed once in IF buffer and incubated with secondary antibodies anti-Rabbit IgG Alexa Fluor Plus 555 (ThermoFisher Scientific, #A32794) in IF buffer supplemented with 10% donkey serum for 6h on a rocker at room temperature. DAPI (Sigma-Aldrich, #MD0015) was added to the organoids and then washed first with water and then with DPBS. Afterwards, organoids were resuspended in ProLong Gold Antifade Mountant (ThermoFisher Scientific, #P36930) and mounted on a glass coverslip. Acquisition was done in a LAS X Leica confocal scanning microscope (Leica) and a 20x Nikon oil-objective. Images were deconvolved and analyzed using ImageJ (NIH).

### Quantification and statistical analysis

Data are presented as means ± SEM or SD as indicated in the Figure legend. Each experiment was performed with a minimum of two biological replicates; exact numbers are depicted in associated Figure legends. Except proteomics and mRNAseq analysis, all statistical analyses were performed using Prism10 software (GraphPad). The statistical tests used and relevant *P* values are mentioned in appropriate Figures legends. *P* values of <0.05 were considered significant.

## Acknowledgments

We are grateful to Roche pool blood donors for valuable support in donating blood. The 360° pRED core facilities for Proteomics (especially for Dr. Angelique Augustin, Sabrina Golling and Justine Fidelin), the Flow cytometry core (especially Dr. Telma Lopes, Dr. Irene Calvo and Dr. Sinduya Krishnarajah) and the IVR-Mouse team at Roche for their invaluable help for this project. We acknowledge the support of the non-profit foundation HTCR, which holds human tissue on trust, making it broadly available for research on an ethical and legal basis. We thanked Marius Franscisco Harter and Giacomo Lazzaroni for their help with the colon organoids culture and Dr. Nicolas Mercado for establishing the collaboration with Fraunhofer. We also acknowledged Biorender (<u>biorender.com</u>) for their imaging collection. Finally, we thank all the CMI department members for their critical feedback and the Roche postdoc fellow (RPF) program for their financial support.

## Extended data Figure Legends

**Extended data Figure 1.**
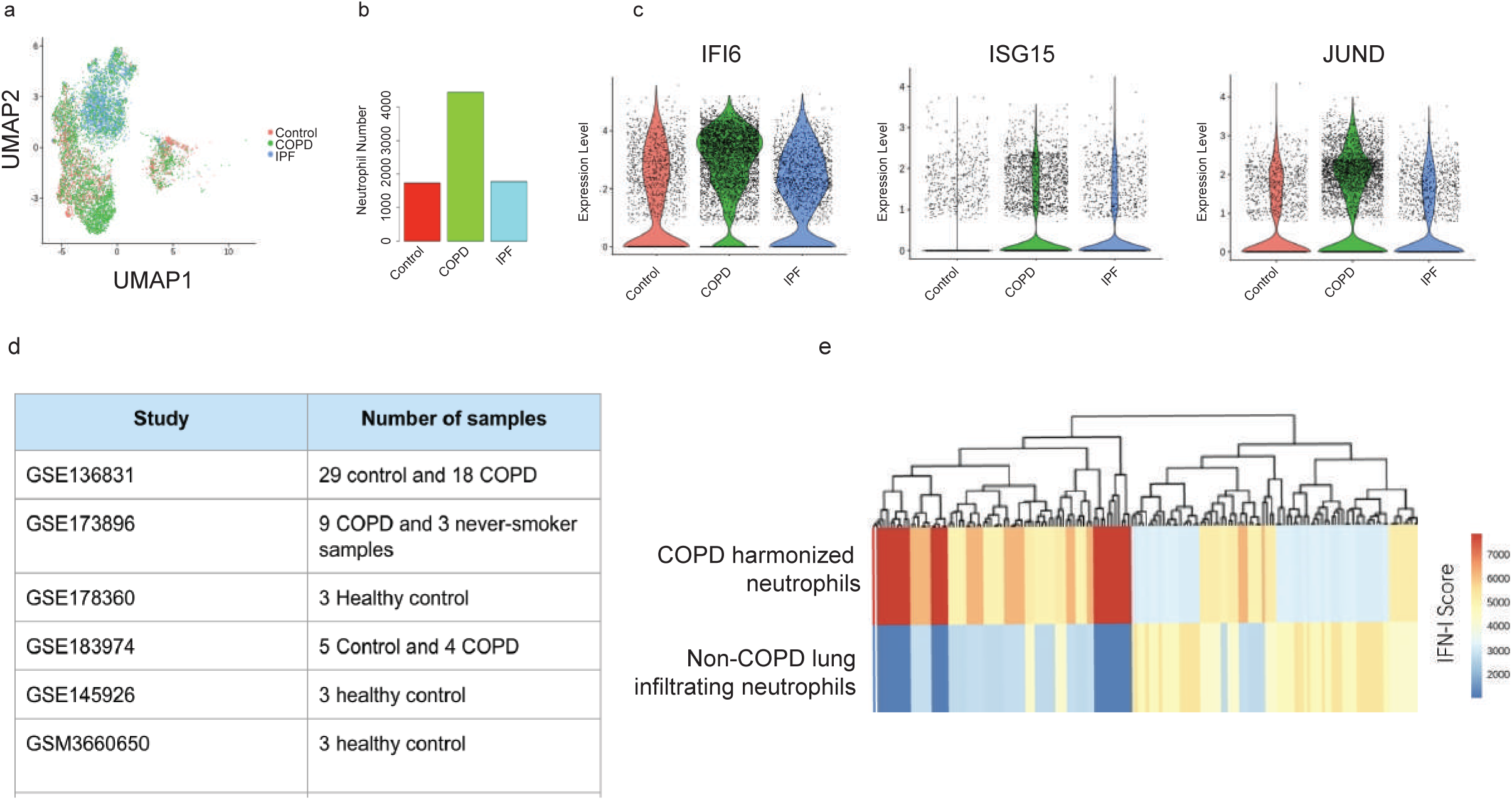
Type I IFN signalling identification in the COPD infiltrating neutrophils. a) UMAP of infiltrating neutrophils UMAP (original NGS from ^65^). The healthy status is depicted by colors and projected on the map. b) Quantification of neutrophils present in the lung of control, COPD or idiopathic pulmonary fibrosis (IPF). c) Violin plots showing the expression levels of type I IFN related genes. d) Table showing the source dataset used to generate the COPD and healthy neutrophil harmonized dataset. e) Heatmap showing the IFN signature, in which red represents high expression and blue represents lower expression. The harmonized dataset was anchored by Celltypist based on the lung plus neutrophil trained models.

**Extended data 2.**
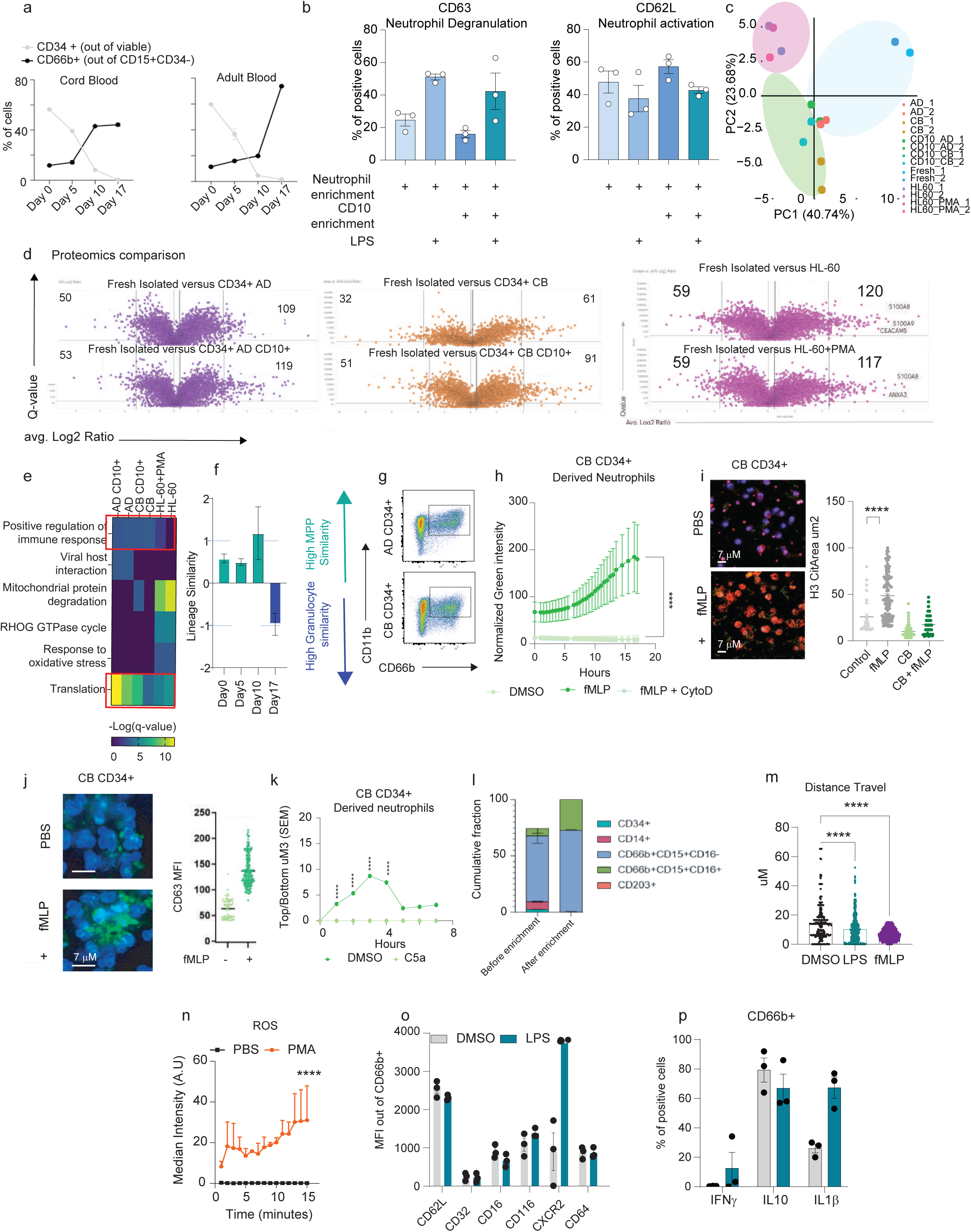
CD34+ in vitro derived neutrophils characterization and functional characterization of the AD-neutrophils. a) Expression levels of CD34 and CD66b positive cells during the neutrophil in vitro differentiation. b) Testing of neutrophil activation upon CD10+ beads selection. LPS was used as control. c) Polar PCA showing the protein similarity between AD, CB derived neutrophils, AD, CB derived neutrophils plus CD10+ further enrichment, HL-60, HL-60 plus PMA and fresh isolated neutrophils. d) Volcano plots showing the identified proteins in comparison to fresh isolated neutrophils. e) Heatmap showing the significant enriched pathways comparing AD, CB derived neutrophils plus CD10+ further enrichment, HL-60, HL-60 plus PMA with fresh isolated neutrophils. f) Bar plots displaying the expression levels of genes associated with progenitor state and neutrophil status across the in vitro differentiation. The plots depict the values obtained from 3 independent differentiation experiments. g) Representative flow cytometry of AD-CD34+ and CB-CD34+ derived neutrophils. h) Quantification of E-Coli ph-Rhodo incorporation across 18hr live imaging. Neutrophils were activated with fMLP and cytochalasin D was used as negative control. Data was normalized by the total number of cells present in the field of view. (n=3 independent experiments; 3 replicates per assay), Statistical analysis was performed by two-way ANOVA followed by Sidak’s multiple comparison. p***<0.001. i) Representative images showing NETs, TRAP formation, of CB-CD34+ derived neutrophils upon fMLP treatment. NETosis is depicted by histone 3 citrullination (H3Cit, red) and nuclei (DAPI, blue). Degranulation is depicted by CD63 (green). Right Panel: Quantification of the TRAP area. Statistical analysis was performed by one-way ANOVA followed by Tukey’s multiple comparison. p****<0.0001. j) Representative images showing neutrophil CB-CD34+ derived neutrophil degranulation depicted by CD63 (Green) upon fMLP treatment. Nuclei is depicted by DAPI (blue). Right panel: Quantification of h. k) Chemoattracted capacity of CB-CD34+ derived neutrophils. l) Flow cytometry characterization of the AD-CD34+ before or after enrichment. m) Quantification of cellular distance travel after LPS, fMLP or DMSO (control). The images were recorded for 4 hours. n) AD CD34+ ROS response upon PMA stimulation. o and p) Flow cytometry characterization of the AD-derived neutrophils upon LPS stimulation. Statistical analysis was performed by one-way ANOVA followed by Tukey’s multiple comparison.

**Extended data Figure 3.**
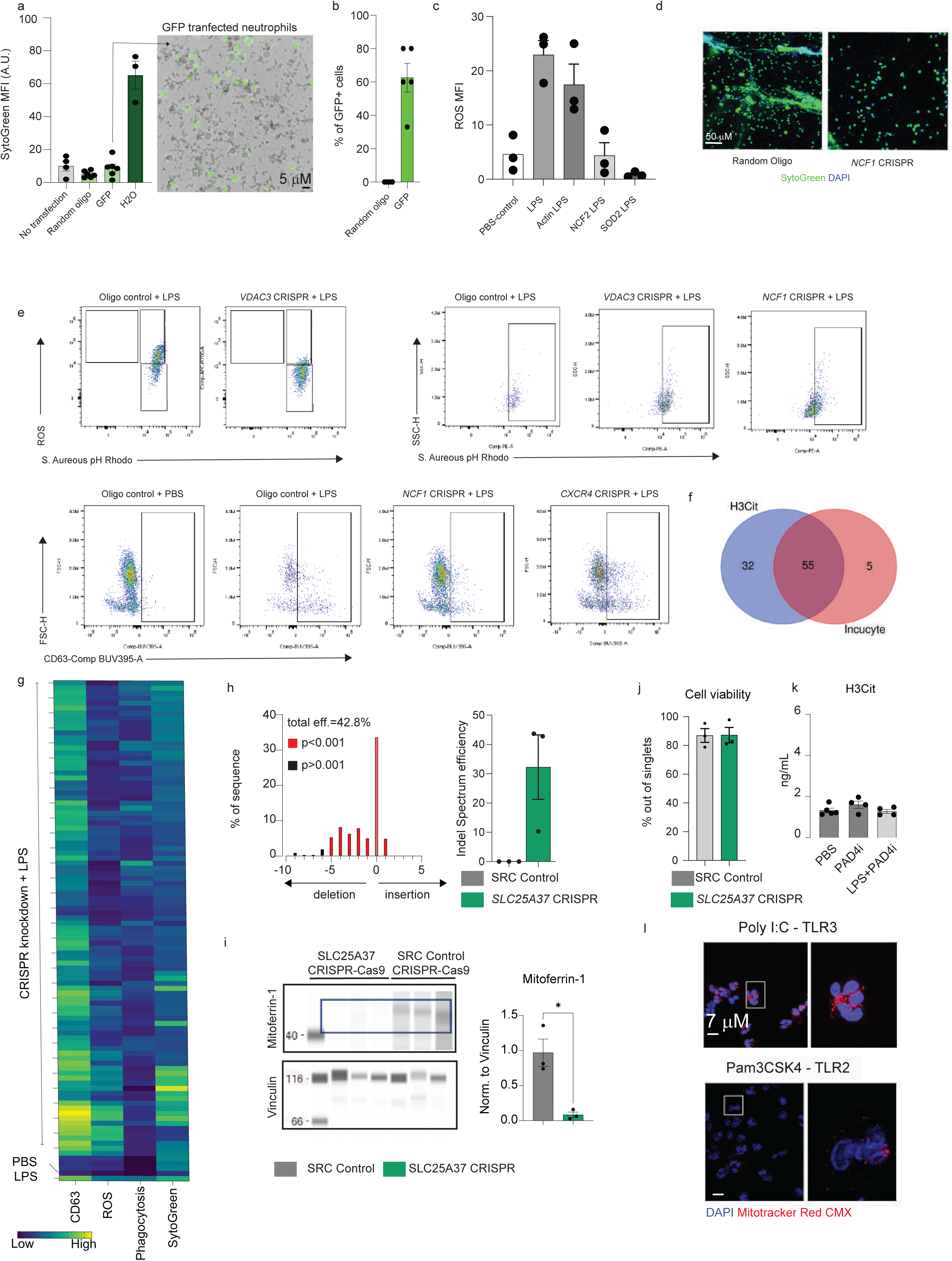
CRISPR-Cas9 screen validation. a) Left panel: Bar-plot showing the fresh neutrophils viability upon transfection. H2O was used as death positive control. Right panel: Representative fluorescence showing the GFP expression after transfection. b) Quantification of GFP+ neutrophils after transfection. c) Bar plots showing the ROS levels upon depletion of *ACTN1, NCF2* and *SOD2.* d) Left panel: Representative immunofluorescence showing the NET formation after *NCF1* depletion. Green represents SytoGreen and nuclei is depicted in blue. Right panel: Quantification of NETosis. e) CRISPR-Cas9 read-out multiplexing examples showing phagocytosis, ROS levels and neutrophil degranulation. LPS was used as positive control. f) Venn diagram showing the common targets found in the imaging (Sytogreen) and H3 citrullination ELISA. g) Heatmap showing the results for phagocytosis, ROS, degranulation and imaging (Sytogreen) for all the 95 targets. h) Left panel: Representative sangar result showing the indels after *SLC25A37* CRISPR editing in fresh isolated neutrophils. Right panel: Quantification of the total efficiency of indels in *SLC25A37* CRISPR edited fresh isolated neutrophils. i) Representative WES blot showing the levels of Mitoferrin-1. Vinculin was used as loading control. j) Neutrophil viability after transfection. k) H3 citrullinate quantification upon LPS stimulation plus or minus PAD4i treatment. l) Representative image showing mitochondrial mass activity (red channel) and nucleus (blue channel) after TLR3 and TLR2 stimulation.

**Extended data Figure 4.**
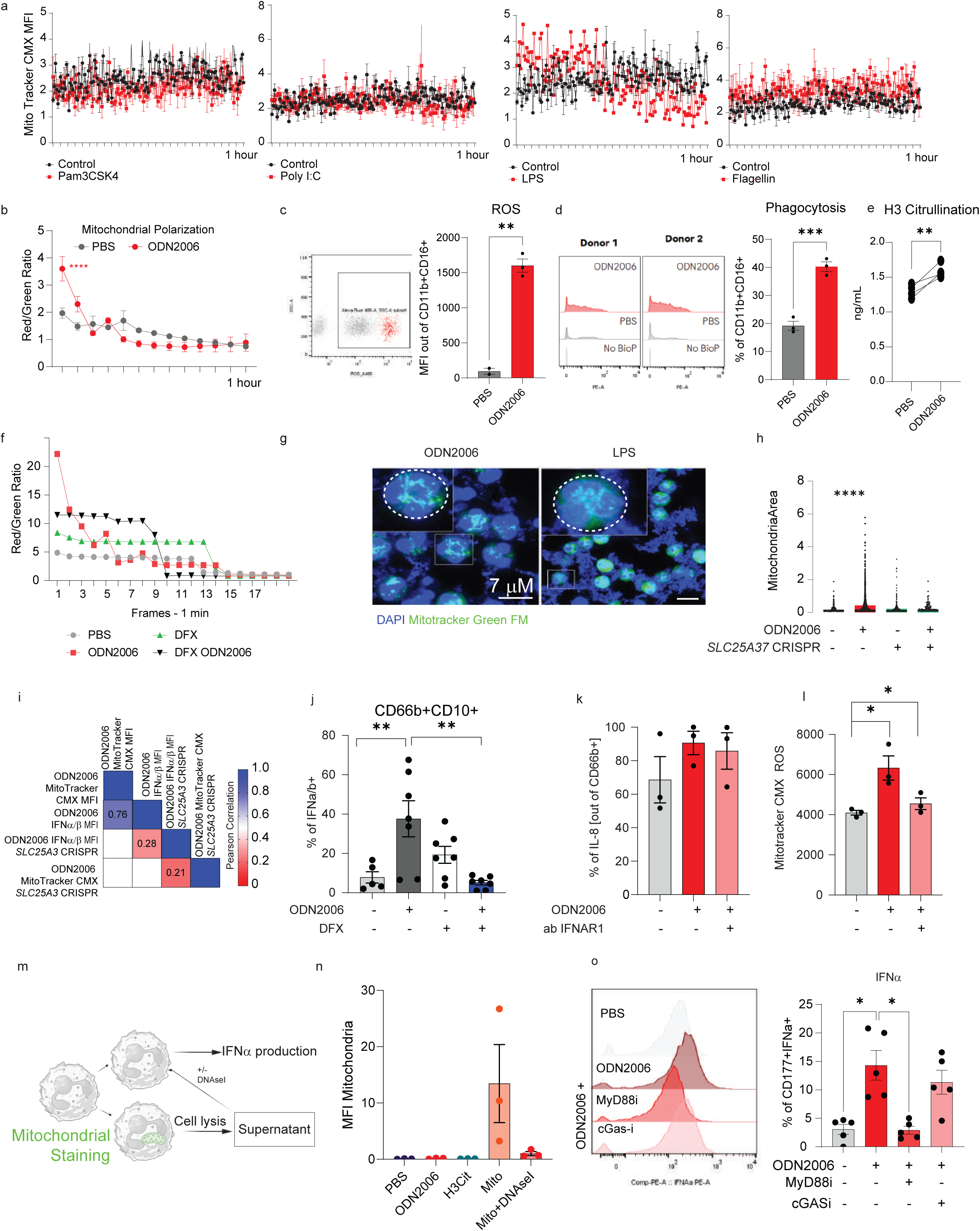
Correlation between mitochondrial activity and type I IFN production. a) Time lapse live imaging quantification displaying the mitochondrial mass activity upon TLRs stimulation. b) Time lapse live imaging quantification showing the mitochondrial polarization based on the red channel to green channel of fresh neutrophils stained by mito probe JC-1. The cells were stimulated with TLR9 agonist (ODN2006). c) ROS levels of fresh isolated neutrophils treated with ODN2006. d) Phagocytosis capacity of fresh isolated neutrophils treated with ODN2006. e) H3 citrullination quantification from the supernatant of fresh isolated neutrophils treated with ODN2006. f) Quantification of mitochondrial polarization based on the red channel to green channel of fresh neutrophils stained by mito probe JC-1. DFX: deferasirox, iron chelator. The TLR9 agonist ODN2006 was used as positive control. g) Representative images showing the mitochondrial morphology (green channel) upon ODN2006 or LPS stimulation. Fresh isolated neutrophils were stained with Mitochondria^Tm^ probe. h) Quantification of the mitochondrial length area. The plugin *MitochondriaAnalyzer* was used. i) Pearce correlation map showing mitochondrial activity and type I IFN production steady state and Mitoferrin1 depletion fresh isolated neutrophils. j) Quantification of type I IFN upon ODN2006 stimulation in the presence of DFX (navy blue) or not. k) Quantification of IL-8 production upon Mitoferrin1 depletion. l) Quantification of Mitochondria CMX upon treatment of fresh isolated neutrophils with IFNAR1 blocking. m) Graphical scheme describing the strategy to acquire autologous mitochondrial DNA (mt-DNA). n) Quantification of the mitochondrial material after co-culture. o) Quantification of type I IFN after stimulation of TLR9 agonist, ODN2006, plus or minus the ubiquitous MyD88 inhibitor (MyD88i) or cGAS-inhibitor, G140.

**Extended data Figure 5. IFNAR1 treatment ameliorates the inflammatory millie in smoked models, but failed to do the same in the DSS-colitis model.**

a) Quantification of neutrophils present in the circulation of IgG, IFNAR1 ab treated or naive mice. b) Median Fluorescence Intensity (MFI) for CD62L in the Neutrophils (Ly6G+CD11b+) and CD63 in c). d) Mitochondrial polarization of Neutrophils (Ly6G+CD11b+) based on JC-1 staining. Statistical analysis was performed by one-way ANOVA followed by Tukey’s multiple comparison. p*=0.05; **p=0.01; ***p=0.001 and p****<0.0001. e) Representative image showing the crypt colon integrity in DSS-Colitis animals treated or not with anti-IFNAR1. f) Diseases activity index characterization. The index is based on body weight, stool/diarrhea and anal bleeding.Statistical analysis was performed by Fisher-test; p*=0.05. g) Percentage of body weight lost during the DSS-Colitis experiment. Statistical analysis was performed by Mann-Whitney test. h) Graphic scheme showing the cigarette smoked COPD model. i) Quantification of infiltrating IFNAR1+ neutrophils naive (CD62L+) and active (CD62L-) in the lung of room air (RA) and cigarette exposed (CS) mice (n=8). j) Quantification of the mitochondrial polarization in the lung infiltrating neutrophils. Statistical analysis was performed by one-way ANOVA followed by Tukey’s multiple comparison. Data variation is presented as ±SEM. k) Left: Representative image showing the Ly6G+ (Neutrophils) infiltration in the lungs of CS animals. Right: Quantification of Ly6G+ cells found per field of view (FOV) from 30x magnification. l) Quantification of the monocytes infiltrating the lungs of cigarette smoked (CS) or room air (RA) mice. The animals were treated with IFNAR1 antibody or IgG isotype control (IgG). m) Neutrophils percentage found in the blood and n) long bone marrow in the same animals as M. o) Barplot showing the MSD quantification of IL-12 levels found in the lung homogenate. Data variation is presented as ±SEM. p) IFNa quantification of the lung homogenate by ELISA. Data variation is presented as ±SEM. Data variation is presented as ±SEM. q) t-SNE map displaying the high dimensionality similarity of the MFI from Ly6G, Ly6C, CD11b, CD63, IFNAR1, JC-1 mito probe (green and red), CD62L, CXCR4, granular complexity (SSC-A) and volume (FSC-A). r) t-SNE map expression of active markers (CD63 and IFNAR1) and naive markers (CD62L and CXC4). The expression level is depicted by the red color.

**Extended data Figure 6. 3D co-culture of colon and alveolar organoids with neutrophils mimics the inflamed milieu found in IBD and COPD**

a) Representative image showing the MCP-1 levels from colon organoids exposed to PBS or inflammatory cocktail. b) Quantification of the MCP-1. c) Permeability assay (TEER) of FITC-Dextran performed in colon monolayer. d) Transwell migration assay of neutrophils towards organoids exposed to PBS or inflammatory cocktail. e) Representative imaging showing the AO and neutrophils depicted with CD66b. f) Quantification of neutrophils observed within the organoids. g) Flow cytometry analysis showing the levels of CXCR2 within the neutrophils (CD66b+). h) Left: Quantification of neutrophils and right: quantification of CXCR2. i) Viability levels of neutrophils in the AO co-culture. j) Levels of caspase 3 cleavage observed in the AO co-cultured with neutrophils. k) Flow cytometry quantification of the infiltrating neutrophils within the AO. l) Viability percentage of the Epcam+ cells after co-culture. m) Representative images showing the IFNa levels. n) Quantification of IFNa found in the CD66b+ cells. Statistical analysis was performed by one-way ANOVA followed by Tukey’s multiple comparison.

